# Siderophores can alter the population dynamics of fungal-bacterial communities by inhibiting specialized metabolism

**DOI:** 10.1101/2025.05.12.653243

**Authors:** Huong T. Pham, Wonyong Kim, Jeongeun Jang, Ji Seon Kim, Young Woon Lim, Ákos T. Kovács, Kyo Bin Kang

**Affiliations:** Research Institute of Pharmaceutical Sciences, College of Pharmacy, Sookmyung Women’s University, Seoul 04310, Korea; Department of Applied Biology, College of Agriculture and Life Sciences, Chonnam National University, Gwangju 61186, Korea; School of Biological Sciences and Institute of Biodiversity, Seoul National University, Seoul 08826, Korea; Institute of Biology, Leiden University, Leiden 2333BE, The Netherlands

**Author notes:** Correspondence: Kyo Bin Kang.

## Abstract

Siderophores are microbial metabolites with a strong affinity for Fe(III) ions, in addition to other roles beyond iron acquisition. In this study, we propose that bacillibactin, a siderophore produced by the bacterium *Bacillus subtilis*, provides an advantage to its producer by inhibiting the specialized metabolism of antagonistic *Penicillium* fungi. Metabolome and transcriptome analyses indicate that iron deficiency caused by bacillibactin plays a critical role in downregulating the specialized metabolism of competing *Penicillium* species. Since many specialized metabolites of *Penicillium* spp. have antibacterial properties, we hypothesize that bacillibactin cross-protects other bacteria during fungal-bacterial community interactions. A tripartite coculture model demonstrated that bacillibactin production influences community dynamics by reducing the growth of specialized metabolite-producing *Penicillium* species, thereby facilitating the colonization of *B. subtilis* and other co-inhabiting bacterial species.

## Introduction

Siderophores are microbial specialized metabolites that strongly chelate Fe(III). To overcome the low solubility and bioavailability of Fe(III), microorganisms secrete siderophores into their environments and subsequently retrieve the Fe(III)-chelated complexes. Beyond their primary role in iron acquisition, recent studies have suggested several additional functions for siderophores^1^. For instance, siderophores can protect the producing organisms from toxic heavy metals^2–4^ and oxidative stress^5–7^, and can also function as signaling molecules that regulate the production of virulence factors^8^.

As key players in bacterial physiology, siderophores are involved in interspecific interactions in various ways. One well-known phenomenon is siderophore exploitation—the heterologous utilization of siderophores produced by other species^9–12^. This dynamic, along with resistance to siderophore exploitation, can drive the evolution of cooperative and competitive behaviors among microbial species^9–13^. Additionally, siderophores have been shown to be inactivated by competitors through biotransformation in both interspecies bacterial and fungal-bacterial interactions^14–16^. Siderophores can also act as sensor-like molecules that detect the presence of competitors^17^, serve as protective agents against oxidative stress during interspecific competition^18^, and provide cooperative cross-protection in polymicrobial communities against antibiotics^19^. Remarkably, a recent study demonstrated that a siderophore can enhance competitive advantage by sensitizing rival bacteria to phage infection, through iron sequestration^20^.

Here, we report that bacillibactin, a catecholate siderophore produced by *Bacillus subtilis*, can inhibit the production of antimicrobial specialized metabolites in competing *Penicillium* species. We initially analyzed the metabolome of 85 *Penicillium* species cocultured with *B. subtilis* IAM 1145 using liquid chromatography–tandem mass spectrometry (LC-MS/MS). We expected that specialized metabolism silenced in *Penicillium* species would be induced during our coculture experiments. However, contrary to our expectations, cocultivation with *B. subtilis* led to a marked decrease in the production of numerous secondary metabolites by *Penicillium* species. We found that bacillibactin produced by *B. subtilis* suppressed the expression of fungal specialized metabolite biosynthetic genes by limiting iron availability to the *Penicillium* species.

Given that *Penicillium* species are known to produce a wide range of antibacterial compounds, we further examined whether bacillibactin influences population dynamics in artificial tripartite fungal– bacterial cocultures. Our results showed that bacillibactin reduces the competitive ability of *Penicillium* species, thereby facilitating the growth of both *B. subtilis* and other co-inhabiting bacterial species. These findings suggest that siderophores may play a pivotal role in shaping the structure of microbial communities.

## Results

### The untargeted metabolomics analysis hypothesized that bacilibactin suppressed specialized metabolism in *Penicillium* species

We analyzed the specialized metabolomes of 85 *Penicillium* species cocultured with *B. subtilis* IAM 1145 using LC-MS/MS, and compared them with those from axenic cultures. Contrary to our initial expectation that *Penicillium*-produced metabolites would increase in the presence of *B. subtilis*, we observed a substantial decrease in the ion intensities of many metabolites in the coculture samples. We examined both the confronting zone—the region on coculture plates where the two organisms physically interact—and the *Penicillium* colonies. In both regions, the detected MS features showed a marked reduction in intensity. Specifically, when comparing extracts from the 85 confronting zones with those from the axenic cultures, 7,380 out of 9,650 MS features exhibited decreased ion intensities (defined as <0.5-fold total sum of ion intensities relative to axenic samples) (Fig. 1a). A similar trend was observed in the *Penicillium* colonies, where 5,874 features showed reduced intensities (Fig. 1b). Although some metabolites were induced in coculture relative to monoculture, these instances were comparatively limited. In the confronting zones, 852 MS features increased (>1.5-fold compared to axenic cultures), and 1,211 features were enriched in the *Penicillium* colonies. This general trend varied in magnitude depending on the *Penicillium* species. *P. brasilianum* exhibited the most dramatic metabolomic shift upon interaction with *B. subtilis*, as shown by base peak ion (BPI) chromatograms displaying a significant reduction in the number of detected metabolites (Fig. 1d). Similar trends were observed for *P. commune*, *P. daejeonium*, *P. virgatum*, *and P. creberum* (Extended Data Fig. 1a–d). However, not all species displayed such extensive changes; for instance, cocultures involving *P. janthinellum* and *P. cairnsense* showed induction of certain metabolites instead (Extended Data Fig. 1e–f).

**Fig. 1.**
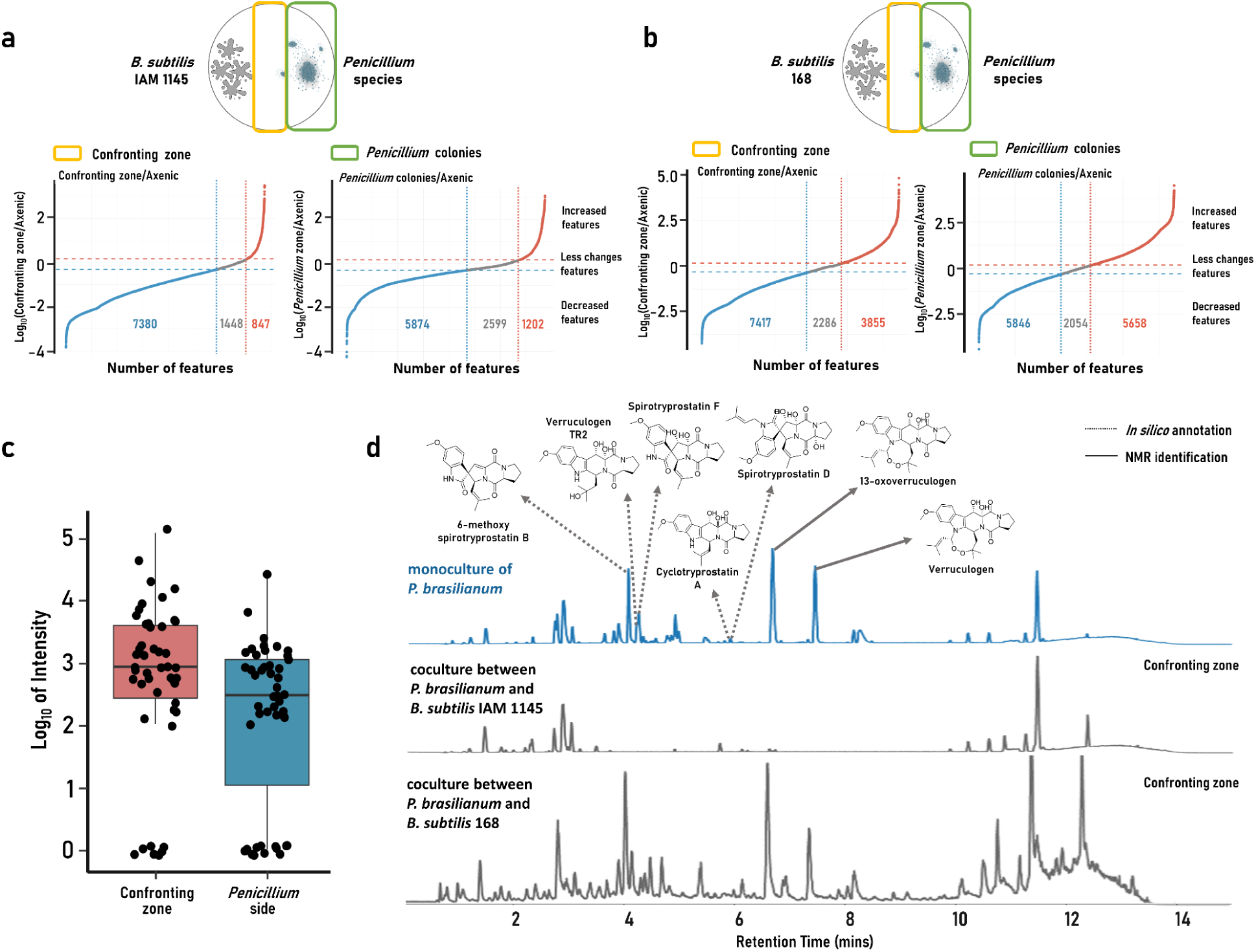
A bacillibactin producer strain *B. subtilis* IAM 1145 reduced specialized metabolite production in *Penicillium* species cocultures, while a non-producer strain *B. subtilis* 168 did not. **a, b**, Changes in MS feature abundances in cocultures of 85 *Penicillium* species with either *B. subtilis* IAM 1145 (**a**) or 168 (**b**), relative to monocultures. Features were categorized based on fold change: decreased (<0.5-fold), unchanged (0.5–1.5-fold), or increased (>1.5-fold). Data from the confronting zone and the *Penicillium* colony regions of the coculture plates were analyzed separately. **c,** MS intensities of the protonated ion of free-form bacillibactin (*m/z* 883.2648, [M+H]⁺) in 47 coculture plates where bacillibactin was detected. Ion intensity was zero in the remaining 38 coculture plates and in the monoculture of *B. subtilis* IAM 1145. **d,** LC-MS base peak ion (BPI) chromatograms of extracts from *P. brasilianum* monoculture and cocultures with either *B. subtilis* IAM 1145 or 168. Peaks corresponding to verruculogen and its analogs are marked with their annotated structures. Verruculogen and 13-oxoverruculogen were confirmed via targeted isolation and NMR analysis.

To characterize the *Penicillium* metabolites that increased or decreased in coculture with *B. subtilis* IAM 1145, we subjected the mass spectrometry data to feature-based molecular networking (FBMN) analysis^21,22^. Due to the limited coverage of the Global Natural Products Social molecular networking (GNPS) spectral library for fungal metabolites, we additionally performed *in silico* annotation of unidentified spectra using Network Annotation Propagation (NAP) with the NPAtlas structural database^23,24^. NAP provided annotations for 3729 features, supplementing the 105 features annotated through GNPS spectral library matching (corresponding to level 2 or 3 annotations according to the Metabolomics Standards Initiative^25^). By carefully evaluating the *in silico*-predicted structures and the corresponding raw MS/MS spectra, we identified candidate annotations likely to be accurate, which allowed us to classify multiple molecular families at the chemical class level (Supplementary Data 1). Metabolites grouped within the same molecular family tended to show coordinated changes—increasing or decreasing together in the coculture—suggesting that the production levels of related congeners shifted in concert. Several molecular families exhibited decreased levels under cocultivation conditions, including alkaloids, anthraquinones, griseofulvins, cyclic peptides, decalins, and terpenoids. In contrast, pyrrocidine-class tetramate alkaloids were enriched during coculture with *B. subtilis* IAM 1145. For a few molecular families, such as the azaphilones, both increases and decreases in individual metabolites were observed (Extended Data Fig. 2).

Upon analyzing the data, we found the enhanced production of bacillibactin in several coculture plates. Bacillibactin was initially annotated via GNPS spectral library matching and subsequently confirmed using a reference standard (Extended Data Fig. 3a). The free form of bacillibactin (unbound to Fe³⁺) was detected above the limit of detection in 47 out of 85 coculture plates (Supplementary Table 1), whereas it was not detected in *B. subtilis* IAM 1145 cultured alone under the same conditions (Fig. 1c). Induction of bacillibactin was observed in several cocultures where fungal specialized metabolism was inhibited, including those involving *P. brasilianum*, *P. virgatum*, *P. creberum*, *P. daejeonium*, and *P. commune*. However, bacillibactin levels and the degree of fungal metabolite inhibition did not show a strict correlation (Extended Data Fig. 3b–f). To determine whether the patterns of metabolite induction and suppression were related to the phylogenetic relationships among *Penicillium* species, we performed a comparative analysis, but no significant association was found (Supplementary Fig. 1).

To assess *Penicillium* specialized metabolite production in the absence of bacillibactin, we cocultured the 85 *Penicillium* species with *B. subtilis* 168 and analyzed the resulting metabolic changes. *B. subtilis* 168 is a domesticated laboratory strain that lacks non-ribosomally synthesized peptides, including bacillibactin, due to a mutation in the gene encoding 4′-phosphopantetheinyl transferase Sfp^26^. In contrast to cocultures with *B. subtilis* IAM 1145, the number of MS features that increased (relative to *Penicillium* axenic cultures) was greater than the number of features that decreased (Fig. 1a and 1b). This pattern represents the opposite trend of what was shown in the *B. subtilis* IAM 1145–*Penicillium* cocultures. These results supported the hypothesis that bacillibactin inhibits specialized metabolism in *Penicillium* species during cocultivation.

### Bacillibactin inhibited the specialized metabolism of *Penicillium*

To further evaluate the hypothesis that bacillibactin influences specialized metabolism in *Penicillium* spp., we selected a representative test species from the 85 screened species for in-depth investigation. *P. brasilianum* was chosen as it exhibited the most pronounced metabolic changes in coculture with *Bacillus subtilis* IAM 1145, while showing minimal response in coculture with *B. subtilis* 168, based on manual inspection of the metabolomics data. The major chromatographic peaks that decreased in coculture with *B. subtilis* IAM 1145 were annotated as verruculogen and related analogs, including 6-methoxy spirotryprostatin B, verruculogen TR-2, spirotryprostatin F, cyclotryprostatin A, spirotryprostatin D, and 13-oxoverruculogen, using *in silico* annotation methods (Fig. 1d; Extended Data Fig. 4). We isolated two major LC-MS peaks and confirmed them as verruculogen and 13-oxoverruculogen by comparing their NMR spectroscopic data with the reference (Supplementary Table 2)^27,28^.

To test our hypothesis that bacillibactin reduces specialized metabolite production in coculture conditions, we compared the effects of *Bacillus subtilis* IAM 1145 and *B. subtilis* 168 on the specialized metabolism of *Penicillium*, and included *B. subtilis* 168 Δ*dhbF*, a mutant strain lacking the *dhbF* gene that encodes a dimodular non-ribosomal peptide synthetase essential for bacillibactin biosynthesis. This mutant was used to prevent potential bacillibactin production in case of spontaneous reconstitution of a functional *sfp* gene. *P. brasilianum* was cocultured with either *B. subtilis* IAM 1145, 168 WT, or 168 Δ*dhbF* for 16 days. Agar media from each coculture were extracted with ethyl acetate (EtOAc) and analyzed by LC-MS/MS. Verruculogen and 13-oxoverruculogen levels were significantly decreased in the coculture with *B. subtilis* IAM 1145, whereas no such clear reduction was shown in cocultures with *B. subtilis* 168 WT or 168 Δ*dhbF* mutant. These results support the role of bacillibactin in suppressing verruculogen and its analogs (Fig. 2a). Similar trends were observed in other *Penicillium* species: MS features annotated as cycloaspeptide A and iso-α-cyclopiazonic acid showed similar reductions in *P. virgatum* cocultures, while features corresponding to averatin and erabulenol showed comparable decreases in cocultures with *P. creberum* (Fig. 2a).

**Fig. 2.**
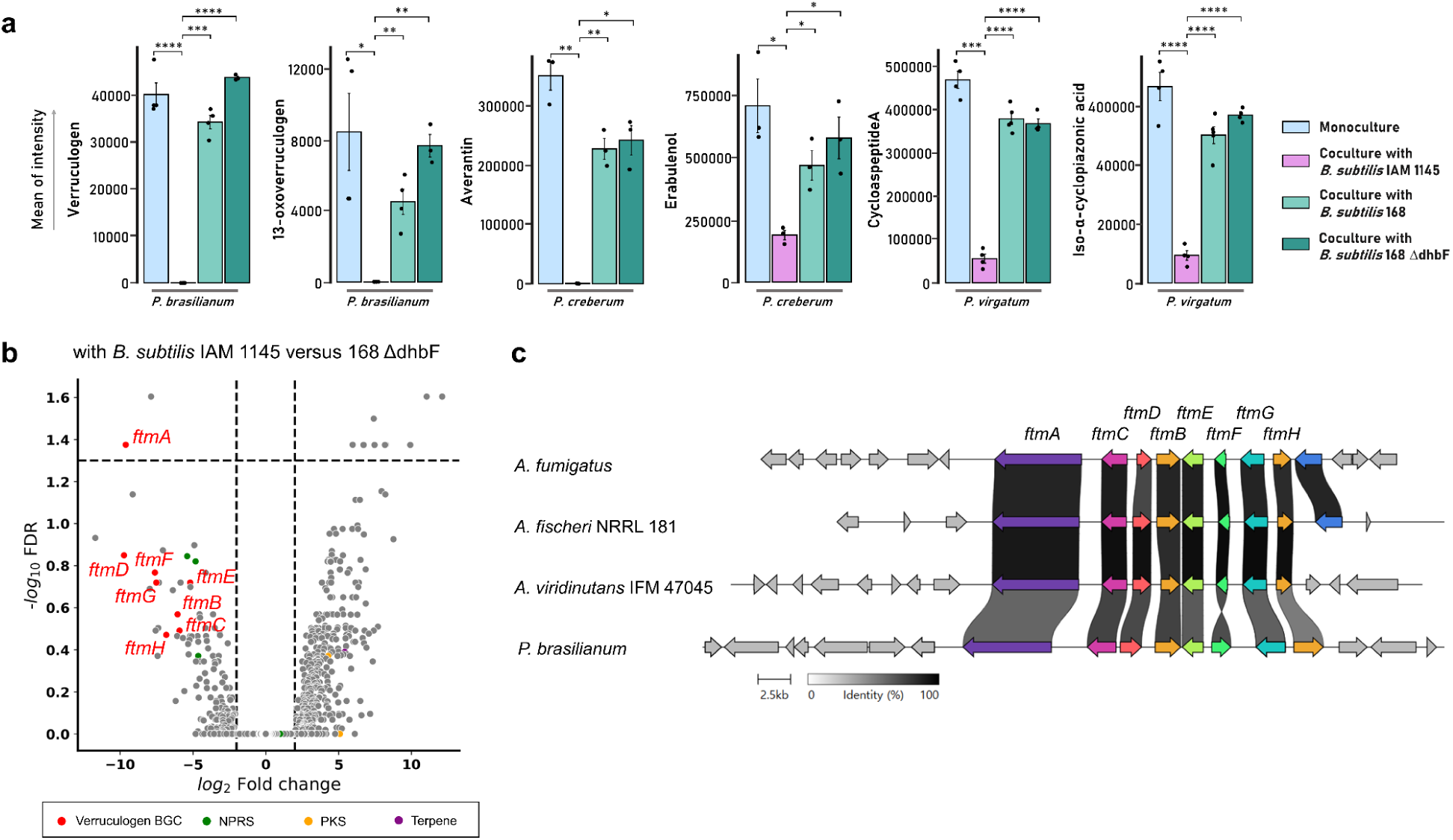
Bacillibactin suppresses specialized metabolite production in *Penicillium* species. **a**, The MS intensities of six compounds produced by three *Penicillium* species were measured in monocultures and cocultures with *B. subtilis* IAM 1145, 168 WT, or 168 Δ*dhbF*. Data are presented as mean ± standard error (SE) from three, four, or five biological replicates for each experiment. Statistical significance is indicated as follows: *p* < 0.05, p < 0.01, *p < 0.001, and **p < 0.0001 by t-test. **b,** RNA-seq analysis of *P. brasilianum*. Fold changes represent the mean expression levels in cocultures with *B. subtilis* IAM 1145 compared to those with *B. subtilis* 168 Δ*dhbF* (three independent biological replicates per condition). Red dots indicate genes in the verruculogen BGC, while green, yellow, and purple dots represent biosynthetic core genes annotated as NRPS, PKS, and terpene synthase, respectively. Grey dots represent other genes. **c,** Comparison of the verruculogen BGCs in three *Aspergillus* species and *P. brasilianum*.

To investigate whether the altered fungal specialized metabolism is due to transcriptional changes, we performed RNA-seq analysis on *P. brasilianum* colonies cocultured with either *B. subtilis* IAM 1145 or *B. subtilis* 168 Δ*dhbF*. We identified 51 biosynthetic gene clusters (BGCs) in the NCBI reference strain *P. brasilianum* MG11, which included 18 polyketide synthases (PKSs), 25 non-ribosomal peptide synthetases (NRPSs) or NRPS-like enzymes, and 8 terpene-related enzymes (Supplementary Data 2). Among these core enzymes of specialized metabolism, the NRPS gene (PMG11_03146), homologous to the brevianamide F synthase FtmA involved in verruculogen biosynthesis in *Aspergillus fumigatus*^29^, was the only differentially downregulated biosynthetic core gene in the coculture with *B. subtilis* IAM 1145 (783-fold downregulated, adj. p = 0.042) (Fig. 2b). Although statistically non-significant, several other NRPS and NRPS-like genes, such as PMG11_01319 (28-fold, adj. p = 0.15), PMG11_07604 (10-fold, adj. p = 0.45), PMG11_08754 (42-fold, adj. p = 0.14), and PMG11_10769 (25-fold, adj. p = 0.43), were also substantially downregulated in the coculture with *B. subtilis* IAM 1145, while a terpene-related gene (PMG11_09714) was highly upregulated (43-fold, adj. p = 0.41) (Fig. 2b; Supplementary Data 2). However, the final products of the NRPS, NRPS-like, or terpene-related genes could not be identified. The verruculogen BGC was highly conserved in *A. fumigatus*, *A. fischeri* NRRL 181, and *A. viridinutans* IFM 47045, comprising 7 tailoring enzyme genes (ftmB–ftmH) (Fig. 2c). The transcriptomic analysis revealed that all these tailoring enzymes were downregulated in the coculture with *B. subtilis* IAM 1145 (Fig. 2b). Therefore, we conclude that the decreased production of verruculogen and its analogs in the presence of bacillibactin-producing *B. subtilis* is attributable to the reduced expression of biosynthetic genes involved in verruculogen production in *P*. *brasilianum*.

### Iron starvation caused by bacillibactin is a cue for *Penicillium* to downregulate specialized metabolism

Since bacillibactin is a siderophore, we hypothesized that iron starvation induced by bacillibactin was the primary factor leading to the reduced production of specialized metabolites by *Penicillium* species. To assess whether bacillibactin caused iron deficiency in the coculture condition, we performed a Chrome Azurol A (CAS)-based assay^30^. CAS is a colorimetric dye that changes from blue in the presence of iron to pink when iron is absent. Among the 85 cocultures, 64 showed iron deficiency around the *Penicillium* colonies, which were characterized by a pink color (Fig. 3a and Supplementary Fig. 2). This group included cocultures of *P. brasilianum*, *P. virgatum*, and *P. creberum*. In 47 of these samples, the presence of free-form bacillibactin was confirmed through LC-MS/MS data (Supplementary Table 1). Bacillibactin was not detected in the LC-MS data of the remaining 21 samples, despite showing signs of iron deficiency. This could be attributed to the relatively low amounts of bacillibactin produced in these cocultures, which may have been below the detection limit of LC-MS. Since Fe^3+^-bacillibactin complexes are difficult to detect in LC-MS due to their instability in acidic conditions^31^, we could not experimentally evaluate this hypothesis.

**Fig. 3.**
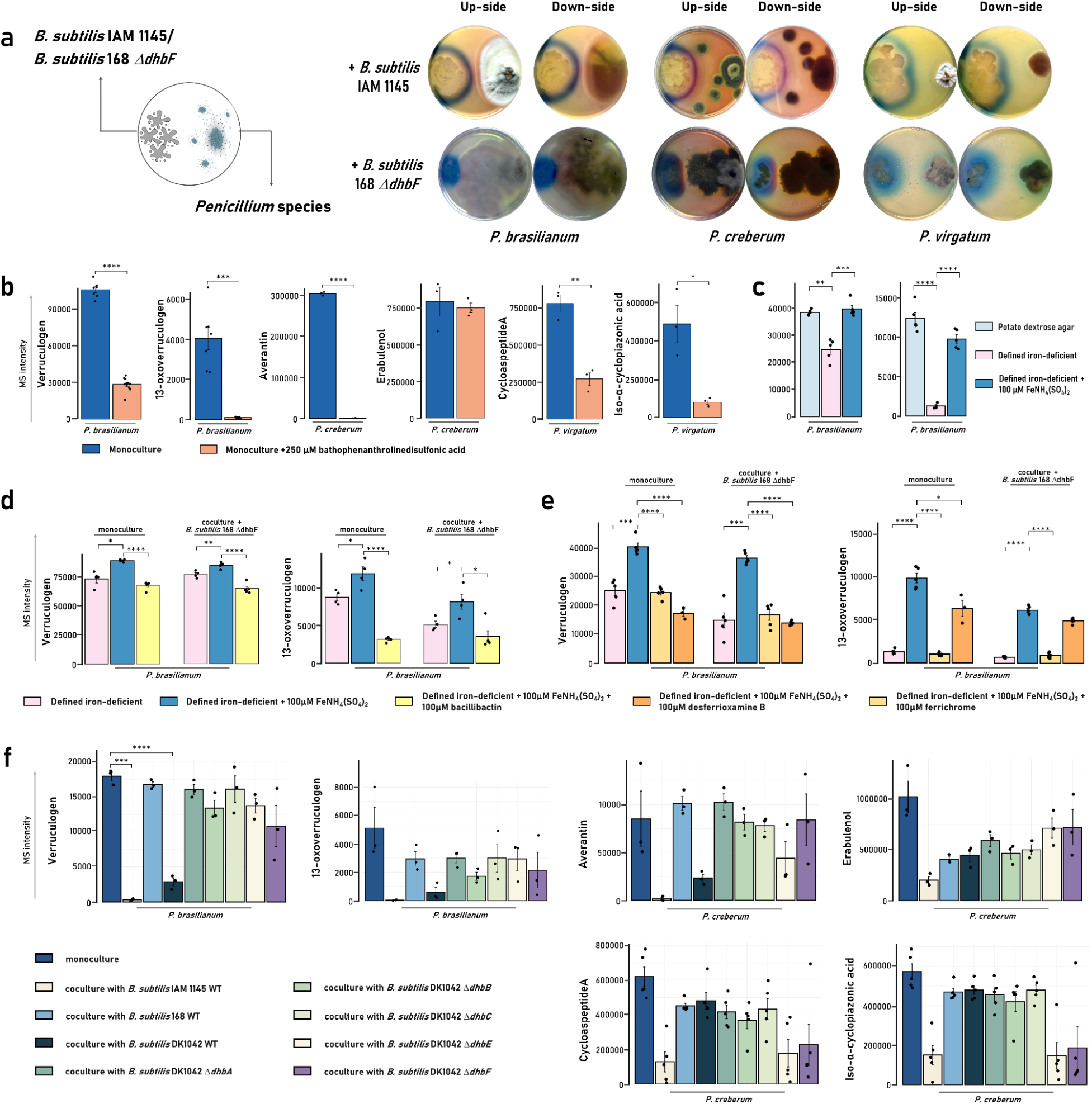
Bacillibactin caused iron deficiency around the *Penicillium* colonies in cocultures, and the iron limitation decreased specialized metabolism. **a**, The Chrome Azurol S assay results of the cocultures between three *Penicillium* species (*P. brasilianum*, *P. creberum*, and *P. virgatum*) and *B. subtilis* IAM 1145 or 168 Δ*dhbF*. Blue color indicates the presence of iron in the media while pink represents the absence. **b**, Comparison of MS intensities of six *Penicillium* metabolites in the extracts of *Penicillium* colonies monocultured in normal PDA medium and iron-depleted PDA medium (by adding 250 μM bathophenanthrolinedisulfonic acid). **c**, MS intensities of verruculogen and 13-oxoverruculogen in the *P. brasilianum* extracts monocultured in the PDA medium or the defined iron-deficient medium designed by us, and the medium supplemented with 100 µM FeNH4(SO4)2. **d**,**e**, MS intensities of verruculogen and 13-oxoverruculogen in the *P. brasilianum* monocultured or cocultured with *B. subtilis* 168 Δ*dhbF* in the defined iron-deficient medium, the medium supplemented with 100 µM FeNH4(SO4)2, and the medium with 100 µM FeNH4(SO4)2 and 100 µM of bacillibactin (**d**), ferrichrome, or desferrioxamine B (**e**). **f**, MS intensities of six *Penicillium* metabolites in three *Penicillium* species, monocultured or cocultured with *B. subtilis* DK1042 WT or Δ*dhbA–F*. Data are presented as mean ± SE of three or five biological replicates for each experiment. \**p* < 0.05, \*\**p* < 0.01, \*\*\**p* < 0.001, and \*\*\*\**p* < 0.0001 by *t*-test.

To confirm that bacillibactin inhibits the specialized metabolism of *Penicillium* species through iron starvation, we cultured the *Penicillium* species in iron-depleted medium and compared metabolite levels with those in the original cultivation medium. When the iron chelator bathophenanthrolinedisulfonic acid was added to remove Fe(II) and Fe(III) from the PDA medium, the levels of verruculogen and 13-oxoverruculogen biosynthesized by *P. brasilianum* decreased, as determined by LC-MS analysis (Fig. 3b). Iron deficiency also led to reduced production of other metabolites, such as cycloaspeptide A, iso-α-cyclopiazonic acid, and averatin. However, the production of erabulenol was not affected by iron restriction, suggesting that bacillibactin may influence erabulenol production in the coculture between *P. creberum* and *B. subtilis* IAM 1145 independently of iron deficiency. Supplementation with 100 µM FeNH4(SO4)2 restored the production of verruculogen and its analogs, indicating that the biosynthesis of verruculogens is closely correlated with the level of bioavailable iron (Fig. 3c). These findings support the hypothesis that the metabolic inhibitory effect of bacillibactin is mediated by iron starvation around *Penicillium* colonies.

We evaluated whether *P. brasilianum* can heterologously utilize bacillibactin. To test this, we added 100 μM ammonium iron(III) sulfate [FeNH₄(SO₄)₂], with or without an equivalent amount of bacillibactin, to a defined iron-deficient medium. We then cultured *P. brasilianum* and measured the production levels of verruculogen and 13-oxoverruculogen. Both compounds showed increased levels under iron-supplemented conditions compared to the control, but their levels decreased in the presence of bacillibactin (Fig. 3d). A similar pattern was observed in a coculture of *P. brasilianum* and *B. subtilis* 168 Δ*dhbF* under the same conditions. This supports the conclusion that bacillibactin, rather than other factors from *B. subtilis*, is responsible for suppressing verruculogen biosynthesis. These results suggest that *P. brasilianum* is unable to utilize the Fe³⁺-bacillibactin complex. Additionally, we tested whether other siderophores could have a similar effect on *Penicillium* specialized metabolism. Two hydroxamate siderophores, ferrichrome and desferrioxamine B, were used in the same experimental setup. Both compounds led to a reduction in the production of verruculogen and 13-oxoverruculogen in *P. brasilianum* (Fig. 3e). These findings suggest that siderophores other than bacillibactin can also inhibit specialized metabolism in fungal competitors that cannot utilize them, likely by inducing iron starvation.

Bacillibactin is biosynthesized from chorismate through a series of reactions. Chorismate is first converted to 2,3-dihydroxybenzoate (2,3-DHB) via enzymes DhbC (isochorismate synthase), DhbB (isochorismate lyase), and DhbA (2,3-dihydro-2,3-DHB dehydrogenase). The NRPS complex DhbEBF then uses 2,3-DHB as a starter unit to elongate a glycine and a threonine unit, followed by cyclization of three monomers to yield bacillibactin^26^. The intermediate 2,3-DHB also has siderophore activity, though its efficacy is lower than that of bacillibactin^32^. We hypothesized that 2,3-DHB could also suppress the specialized metabolism of competitor fungi. However, our results with *B. subtilis* 168 indicated that 2,3-DHB itself was unlikely to have this effect, since *B. subtilis* 168 can produce 2,3-DHB.

To test this hypothesis and gather more direct evidence on the influence of bacillibactin, we used *B. subtilis* DK1042, a competent strain derived from *B. subtilis* NCIB3610 that can produce bacillibactin^33^, and its single gene disruption mutants (Δ*dhbA*, Δ*dhbB*, Δ*dhbC,* Δ*dhbE*, and Δ*dhbF*)^34^. In coculture experiments, *B. subtilis* DK1042 wild-type (WT) suppressed the production of verruculogen and 13-oxoverruculogen in *P. brasilianum*, while all of the Δ*dhbA*–*F* mutants did not affect metabolite production (Fig. 3f). This pattern was similar to the results observed with *B. subtilis* IAM 1145 and 168 cocultures, suggesting that 2,3-DHB plays a very weak or no role in inhibiting specialized metabolism, at least at the concentrations produced in our coculture experiments.

In *P. creberum*, averantin showed a suppression pattern similar to verruculogen while erabulenol was consistently reduced across all cocultures, regardless of the presence or absence of bacillibactin. This suggests that the suppression of erabulenol production may be due to other factors, consistent with the findings under iron-restricted conditions (Fig. 3b). For *P. virgatum*, neither cycloaspeptide A nor iso-α-cyclopiazonic acid showed significant decreases in the cocultures with *B. subtilis* DK1042 WT or its mutants. Bacillibactin was not detected in the LC-MS/MS data from the coculture of *P. virgatum* with *B. subtilis* DK1042 WT and its mutants (Supplementary Fig. 3), indicating that *P. virgatum* may not induce bacillibactin biosynthesis in *B. subtilis* DK1042, and consequently, no inhibitory effect was observed.

### Siderophore can cross-protect other bacteria in community coexistence

*Penicillium* species are well-known for their ability to produce a variety of antibiotics. Several secondary metabolites whose production was suppressed in cocultures with *B. subtilis* IAM 1145 are known to exhibit antibacterial properties^35–38^. Thus, we hypothesized that the bacillibactin-mediated suppression of fungal specialized metabolite production could facilitate the survival of other bacteria within fungal-bacterial communities. To test this hypothesis, we designed a tripartite coculture experiment consisting of one *Penicillium* species, one *B. subtilis* strain, and an additional bacterial species. In this setup, *Escherichia coli* or *Pseudomonas putida,* as the third partner in the coculture,s was inoculated with either *B. subtilis* IAM 1145, 168, or DK1042 (WT or Δ*dhbF* mutants) (Fig. 4a).

**Fig. 4.**
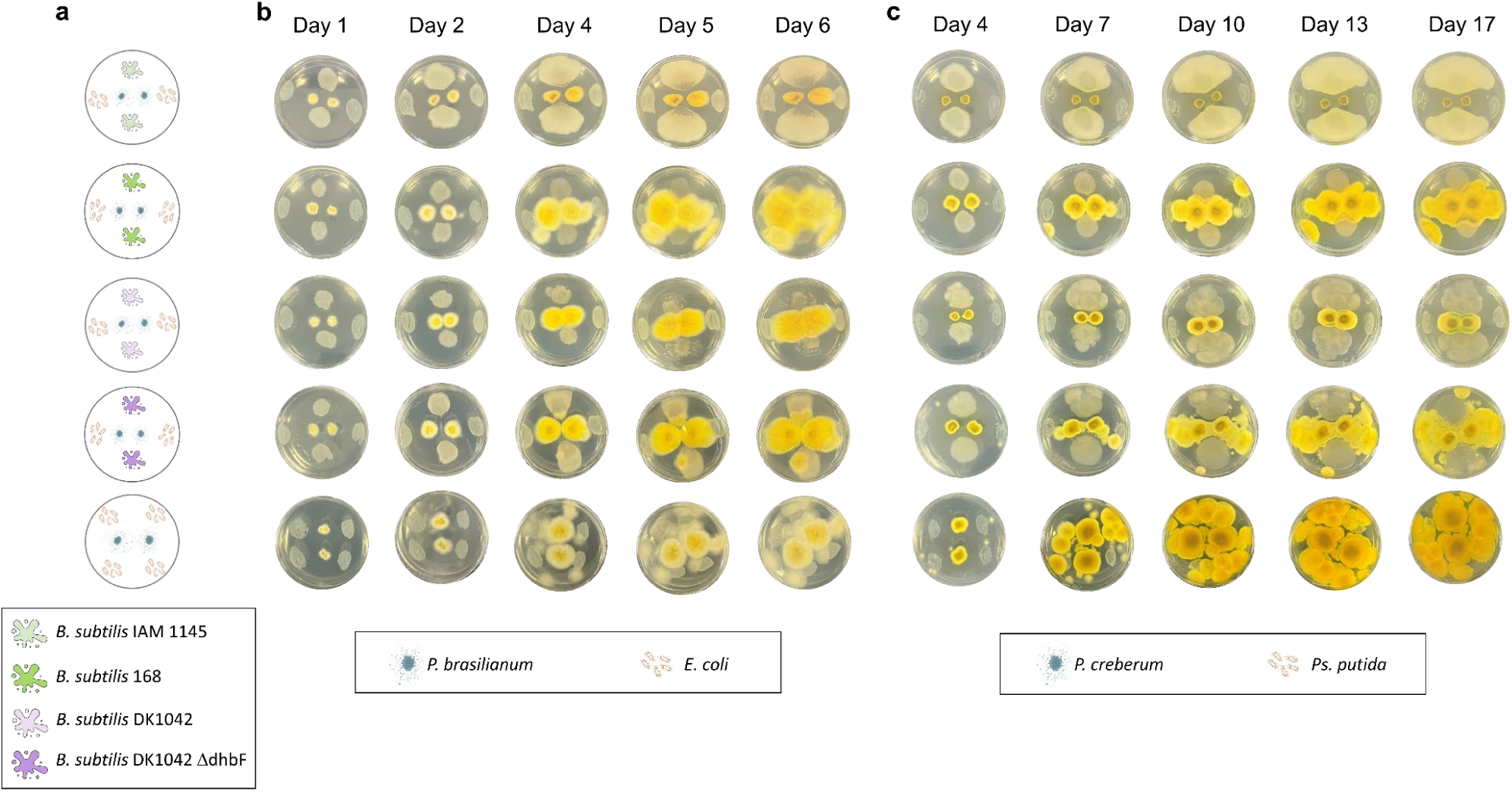
Bacillibactin alters the dominant member of tripartite cocultures of *Penicillium* spp.-*B. subtilis*-*E. coli* or *Ps. putida*. **a**, The plate design for the tripartite cocultures. **b,c** Time-dependent observations of the plates. *P. brasilianum* was cultured with *E. coli* as the third bacterial partner in **b**, while *P. creberum* was cultured with *Ps. putida* in **c**. The dynamics of colony growth were monitored over time.

The presence of bacillibactin-producing *B. subtilis* strains influenced the microbial species that became dominant in the tripartite cocultures. When either *B. subtilis* 168, DK1042 Δ*dhbF*, or no *B. subtilis* was included, the *Penicillium* species dominated, overgrowing all bacterial colonies on the agar medium. However, in cocultures containing the bacillibactin-producing *B. subtilis* IAM 1145 or DK1042 WT, a different dynamic emerged. The growth of the *Penicillium* colonies was significantly restricted, and both *B. subtilis* IAM 1145 or DK1042 WT and the other bacterial partner (either *E. coli* or *Ps. putida*) survived on the agar medium. This survival was likely due to the reduced production of antibacterial metabolites by the *Penicillium* species, which otherwise would have benefited the fungal fitness in competitive environments. This pattern was particularly evident in a time-dependent manner in the tripartite cocultures of *P. brasilianum* - *B. subtilis* - *E. coli* (Fig. 4b) and *P. creberum* - *B. subtilis* - *Ps. putida* (Fig. 4c). Similar trends were also observed in other combinations, such as *P. brasilianum* - *B. subtilis* - *Ps. putida* and *P. creberum* - *B. subtilis* - *E. coli* (Extended Data Fig. 5). These results suggest that bacillibactin cross-protects other bacterial species in microbial communities by inhibiting the specialized metabolism of antagonistic fungi.

## Discussion

Iron is a crucial nutrient for various biological processes essential for life, including respiration, DNA synthesis, and translation. Given its importance, iron deficiency leads to significant physiological and metabolic changes in microorganisms. It is well established that iron deficiency triggers the biosynthesis of siderophores in many bacterial species^39^. However, the impact of iron-limited environments on microbial communities has been less frequently explored. In coculture settings, the biosynthesis of the antibiotic metabolite actinorhodin by *Streptomyces coelicolor* is enhanced due to iron competition mediated by myxochelin, a siderophore produced by *Myxococcus xanthus*^40^. This competition also induces the expression of several BGCs for specialized metabolites in *Streptomyces*. Conversely, the suppression of specialized metabolism due to iron restriction is a relatively rare phenomenon. For instance, Lyng and colleagues demonstrated that bacillibactin inhibits the growth of *Pseudomonas marginalis* by repressing the transcription of the histidine kinase-encoding gene *gacS* and downregulating the expression of biosynthetic genes responsible for producing specialized metabolites like pyoverdine and viscosin^34^.

Our study demonstrates that bacillibactin suppresses a range of specialized metabolites in multiple *Penicillium* species. Moreover, the iron starvation caused by bacillibactin appears to be the primary factor responsible for inhibiting the specialized metabolism of *Penicillium* species. This suppression of specialized metabolites was also replicated in axenic cultures of *Penicillium* species cultured in chemically defined, iron-deficient media. While the precise molecular mechanism of this metabolic suppression remains unclear, we hypothesize that it may be a result of metabolic adaptation. Previous studies on *Saccharomyces cerevisiae* have shown that cellular metabolism can adjust to optimize the utilization of limited iron^41,42^. Many enzymes involved in specialized metabolism, such as cytochrome P450s, require iron as a cofactor. Thus, under conditions of iron limitation, fungi may suppress the expression of specialized metabolic genes unless the metabolites contribute to iron acquisition. The suppression of specialized metabolism in competing microorganisms could represent an auxiliary function of siderophores. Indeed, manipulating iron availability can significantly influence community dynamics^43^. For example, the volatile trimethylamine increases the environmental pH, depleting iron availability, and induces siderophore production in *Streptomyces venezuelae*. This excessive depletion of iron by *S. venezuelae* negatively affects the growth of other soil bacteria and fungi.

Our experiments with ferrichrome and desferrioxamine B indicated that other siderophores can also inhibit the specialized metabolism of competing microorganisms. Given that iron deficiency is the mechanism behind this effect, it was not entirely surprising, as all siderophores can induce iron starvation in the environment. However, it was particularly interesting that ferrichrome suppressed verruculogen biosynthesis, as ferrichrome is a siderophore known to be produced by various fungal species, including *Ustilago sphaerogena*, *Aspergillus quadricinctus*, and *Penicillium chrysogenum*^44–46^. This result suggests that siderophore-mediated suppression of specialized metabolism could occur in fungal-fungal interactions. However, additional experimental evidence is needed to validate this hypothesis further.

Using a tripartite coculture experimental setup, we demonstrated that iron depletion through siderophore production plays a key role in determining the dominant species within microbial communities. Predicting population dynamics in complex microbial environments is challenging, as interactions may vary in different settings, such as pairwise or single-species cultures. Chang and colleagues found that 71.5% of pairwise bipartite co-cultures from 12 stable enrichment communities exhibited competitive exclusion^47^. While our tripartite co-culture models were artificially assembled, they revealed that a single metabolite—specifically a siderophore—can significantly influence community composition and interactions between its members. A similar phenomenon was observed in the THOR three-species model community, composed of *Flavobacterium johnsoniae* UW101, *Pseudomonas koreensis* CI12, and *Bacillus cereus* UW854^48^. In that work, metatranscriptomic and metabolomic analyses of monocultures, bipartite, and tripartite co-cultures showed that biosynthetic gene regulation and metabolite production were dramatically altered by community composition. The authors highlighted how a single metabolite from *Ps. koreensis*, the antimicrobial koreenceine, strongly influenced the regulation of specialized metabolism in the other species, although the exact mechanism of this interaction remains unclear. Siderophore-mediated microbial community differentiation has also been demonstrated in natural rhizosphere microbiomes^49^. Gu and colleagues showed that multiple siderophore-producing bacteria protect tomato plants from the pathogenic bacterium *Ralstonia solanacearum* through iron competition. This growth inhibition by siderophores occurred only when *R. solanacearum* lacked receptors for heterologous siderophore uptake, as observed in other bacterial species^10,50^. In our study, a similar phenomenon was observed in fungal-bacterial interactions, where *Penicillium* species could not utilize bacillibactin, resulting in inhibited growth and suppression of their specialized metabolism. It remains unclear whether *R. solanacearum* is affected by a comparable mechanism in the rhizosphere. While the generality of our findings regarding bacillibactin-mediated interactions between *Penicillium* and *B. subtilis* is still to be confirmed in other microbial communities, the widespread presence of siderophores in nature and their role in iron homeostasis suggest that siderophore-mediated suppression of specialized metabolism could play a significant role in shaping microbial community structures more than previously anticipated.

## Methods

### Microorganisms, media, and cultivation

All *Penicillium* species used in this study (Supplementary Table 3) were obtained from Korea Marine Fungi Resource Bank and maintained on potato dextrose agar (PDA) (Kisan Bio Co., Ltd.) at 25 °C. *B. subtilis* IAM 1145 and 168, *E. coli* NCTC 9001, and *Ps. putida* A.3.12 were obtained from the Korean Collection for Type Cultures, Korea Research Institute of Bioscience and Biotechnology. *B. subtilis* 168 Δ*dhbF* (BKE31960) mutant strain was sourced from the *B. subtilis* gene deletion library^51^ and provided by the National BioResource Project (National Institute of Genetics, Japan). *B. subtilis* DK1042 WT and its Δ*dhbA–F* derivatives were previously described^33,34^. All *B. subtilis* strains were maintained on PDA medium at 25 °C. *E. coli* and *Ps. putida* were maintained in lysogeny broth (LB) medium and stored in small aliquots at − 80 °C. To induce iron deficiency in PDA medium, 250 μM of bathophenanthrolinedisulfonic acid disodium salt hydrate (Sigma-Aldrich) was added. For the chrome azurol S (CAS) assay^30^, CAS-PDA medium was prepared by mixing 60.5 mg of CAS (dissolved in 50 mL H2O) with 1 mM FeCl3·6H2O in 10 mM HCl (10 mL), and combining this with 750 mL of PDA medium, adjusting the final volumen to 1 L with distilled water. The composition of the defined iron-deficient medium we designed is provided in Supplementary Table 4.

### Coculture of *Penicillium* species with *B. subtilis*

Agar plugs (5 mm × 5 mm) of each *Penicillium* species were excised from 7-day-old culture plates and placed on one side of fresh PDA plates. Simultaneously, 5–10 colonies of *B. subtilis* were transferred using an inoculation loop to the opposite side of the plate, covering approximately a 5 mm × 5 mm area. The cocultures were incubated for 16 days at 25 °C. After the incubation period, the solid agar containing the colonies was dissected from the coculture plates, cut into small pieces, and extracted with 40 mL of ethyl acetate (EtOAc). The organic extracts were dried and re-dissolved in 50% aqueous methanol to a final concentration of 1 mg/mL. The samples were then filtered through PTFE syringe filters (Altoss) and subjected to LC-MS/MS analysis. For comparison, monocultures of all 85 *Penicillium* species and *B. subtilis* were grown under identical conditions and processed using the same extraction and sample preparation procedures.

### LC-MS/MS data acquisition

LC-MS/MS analyses were performed using a Waters ACQUITY UPLC system (Waters Corp.) coupled to a Waters VION IMS QTOF mass spectrometer (Waters MS Technologies), equipped with a standard electrospray ionization (ESI) source. Each sample (2 μL) was injected onto an ACQUITY UPLC BEH C18 column (2.1 × 100 mm, 1.7 μm particle size), operated at a flow rate of 0.3 mL/min. The mobile phase consisted of 0.1% formic acid in water (solvent A) and acetonitrile (solvent B). A linear gradient from 10% to 100% solvent B was applied over 0–12 min, followed by a 3 min washout at 100% B, and a 3 min re-equilibration at 10% B. MS/MS spectra were acquired in positive ion mode using MS^E^ data-independent acquisition. Low-energy scans (6 eV) were used for precursor ion detection, while high-energy scans (20–40 eV) enabled fragmentation of precursor ions for structural elucidation.

### LC-MS/MS data analysis for cocultures between 85 *Penicillium* species and *B. subtilis*

LC-MS/MS raw data were processed using MS-DIAL version 4.60 for feature detection and MS/MS spectral deconvolution^52^. Processing parameters included a minimum peak detection height of 10,000, an *m/z* tolerance of 0.01, and a retention time (tR) tolerance of 0.05 min. The resulting MS feature table and MS/MS spectra were uploaded to the Global Natural Products Social (GNPS) web platform (https://gnps.ucsd.edu) for Feature-Based Molecular Networking (FBMN)^22^. For *in silico* metabolite annotation, the Network Annotation Propagation (NAP) workflow^53^ was employed using the NPAtlas structural database^23,24^, with the following parameters: the top 10 candidate structures for consensus scoring and mass candidate searches within 5 ppm.

The GNPS molecular networking data are available at: *Penicillium* species *- B. subtilis* IAM 1145 cocultures: https://gnps.ucsd.edu/ProteoSAFe/status.jsp?task=8f16183a659e4ecea3ae9d41a240eb7d.

The NAP *in silico* annotation result for *Penicillium* spp. *- B. subtilis* IAM 1145 cocultures: http://proteomics2.ucsd.edu/ProteoSAFe/status.jsp?task=20f945fad7d4473ba8c5cf200e5cbbeb.

*Penicillium* species *- B. subtilis* 168 cocultures: https://gnps.ucsd.edu/ProteoSAFe/status.jsp?task=6ec7c2e3c2cd4229bd8a4a34a65f6d3e.

## Isolation and structural determination of verruculogen and 13-oxoverruculogen

*P. brasilianum* was cultured on 169 PDA plates (60 × 15 mm) for 16 days at 25 °C. The entire agar medium, including fungal colonies, was dissected and extracted with EtOAc (4 × 2 L), yielding 1.1 g of crude extract. This extract was fractionated into seven subfractions (P1–P7) using a Waters 600 preparative HPLC system equipped with a YMC Actus Hydrosphere C18 column (250 × 20.0 mm, 5 μm, YMC Co., Ltd.), applying a linear gradient of MeCN–H₂O (12 mL/min, 70:30 to 10:90 over 30 min). Fraction P6 was further purified by preparative HPLC using a Spursil C18 EP column (250 × 10.0 mm, 5 μm, Dikma Technologies Inc.) under isocratic elution (4 mL/min, MeCN–H₂O, 35:65 for 28 min), yielding pure verruculogen (1.5 mg, tR = 15.5 min) and five subfractions (P6a–P6e). Subfraction P6a was further purified under isocratic conditions (MeCN–H₂O, 42:58, 4 mL/min, 25 min) using the same column to yield 13-oxoverruculogen (1.2 mg, tR = 7.7 min). Nuclear magnetic resonance (NMR) spectra were recorded using a Bruker Avance III HD 500 MHz spectrometer (Bruker Daltonics) at the Chronic and Metabolic Diseases Research Center, Sookmyung Women’s University. NMR data are in Supplementary Table 2.

### RNAseq analysis on *P. brasilianum* cocultured with *B. subtilis* either IAM 1145 or 168 Δ*dhbF*

Total RNA was extracted from *P. brasilianum* colonies after 16 days of cocultivation with either *B. subtilis* IAM 1145 or the siderophore-deficient *B. subtilis* 168 Δ*dhbF* mutant, using the TRIzol protocol^54^. RNA samples were treated with the RNase-Free DNase I Set (Qiagen) and purified using the RNA Clean & Concentrator-25 kit (Zymo Research), following the manufacturer’s instructions. RNA quality was assessed using an Agilent 2100 Bioanalyzer (Agilent Technologies), with RNA integrity numbers (RIN) ranging from 9.1 to 10.0 across all samples. Sequencing was performed on the Illumina NovaSeq X platform using paired-end 100 bp reads, generating approximately 6 Gb of data per sample. Adapter trimming and quality filtering were conducted using Trim Galore (https://github.com/FelixKrueger/TrimGalore). High-quality reads were aligned to the *P. brasilianum* MG11 reference genome (GenBank accession: GCA_001048715.1) using HISAT2^55^, allowing relaxed mismatch penalties (--mp 3,1) to account for strain differences. Default parameters were used otherwise. On average, 93% of the reads mapped to genic regions in the reference annotation. Gene expression levels were calculated in counts per million (CPM) and normalized using the trimmed mean of M-values (TMM) method implemented in the R package *edgeR*. Genes with CPM > 1 in at least three samples were retained for downstream analysis to eliminate low-abundance transcripts. Differentially expressed (DE) genes were identified based on a >4-fold change in expression and a false discovery rate (FDR) < 0.05. A volcano plot illustrating the DE genes was generated using the *Seaborn* Python visualization library^56^.

### Tripartite coculture of Penicillium spp.-B. subtilis-E. coli or Ps. putida

For tripartite coculture assays, two 5 mm × 5 mm agar plugs of *P. brasilianum*, *P. creberum*, or *P. virgatum* were excised from 7-day-old PDA cultures and placed centrally on a fresh PDA plate. Simultaneously, 5–10 colonies of *B. subtilis* (IAM 1145, 168, DK1042 WT, or DK1042 Δ*dhbF* mutant) were inoculated on both sides of the plate, each occupying an area approximately 5 mm × 5 mm, using a sterile inoculation loop. To introduce the third microbial partner, 1.0 µL of either *E. coli* (1 × 10³–1.5 × 10³ CFU) or *P. putida* (70–90 CFU) was directly inoculated onto the PDA plate from freshly thawed stocks. The specific inoculation positions for each organism are illustrated in Fig. 4a. Plates were incubated at 25 °C, and photographs were taken at time points determined by the relative growth rates of the constituent strains.

## Data availability

All raw and processed LC-MS/MS data are publicly available at the MassIVE repository by the University of California, San Diego Center for Computational Mass Spectrometry website (https://massive.ucsd.edu) under the accession number MSV000097880. Raw sequence reads acquired from the RNAseq analysis have been deposited at the NCBI Sequence Read Archive under BioProject PRJNA1146190.

## Code availability

The code for data cleaning is available at https://github.com/huong168/Siderophores_can_alter_the_population_dynamics_of_fungal_bacterial_communities

## Acknowledgments

This research was supported by the National Research Foundation of Korea (NRF) grants funded by the Ministry of Science and ICT (2020R1C1C1004046, RS-2025-00523337, RS-2022-NR068419, and RS-2022-NR070845) and a Korea Basic Science Institute (National Research Facilities and Equipment Center) grant funded by the Ministry of Education (RS-2024-00436674). This work was also funded by the Marine Fishery Bio-resources Center (2025) of the National Marine Biodiversity Institute of Korea (MABIK). During the preparation of this work, the authors H.T.P. and K.B.K. used ChatGPT to improve the clarity and readability of the originally submitted manuscript. After using this tool, all authors reviewed and edited the content as needed and take full responsibility for the content.

## Author contributions

Conceptualization: H.T.P. and K.B.K.; methodology: H.T.P., W.K., A.T.K., and K.B.K.; investigation: H.T.P., W.K., J.J., and J.S.K.; resourses: A.T.K. and Y.W.L.; visualization: H.T.P. and W.K.; funding acquisition: K.B.K.; supervision: A.T.K. and K.B.K.; writing–original draft: H.T.P. and K.B.K.; writing–review and editing: A.T.K., Y.W.L., and K.B.K.; All authors reviewed and agreed to the published version of the manuscript.

---

**Extended Data Fig. 1.**
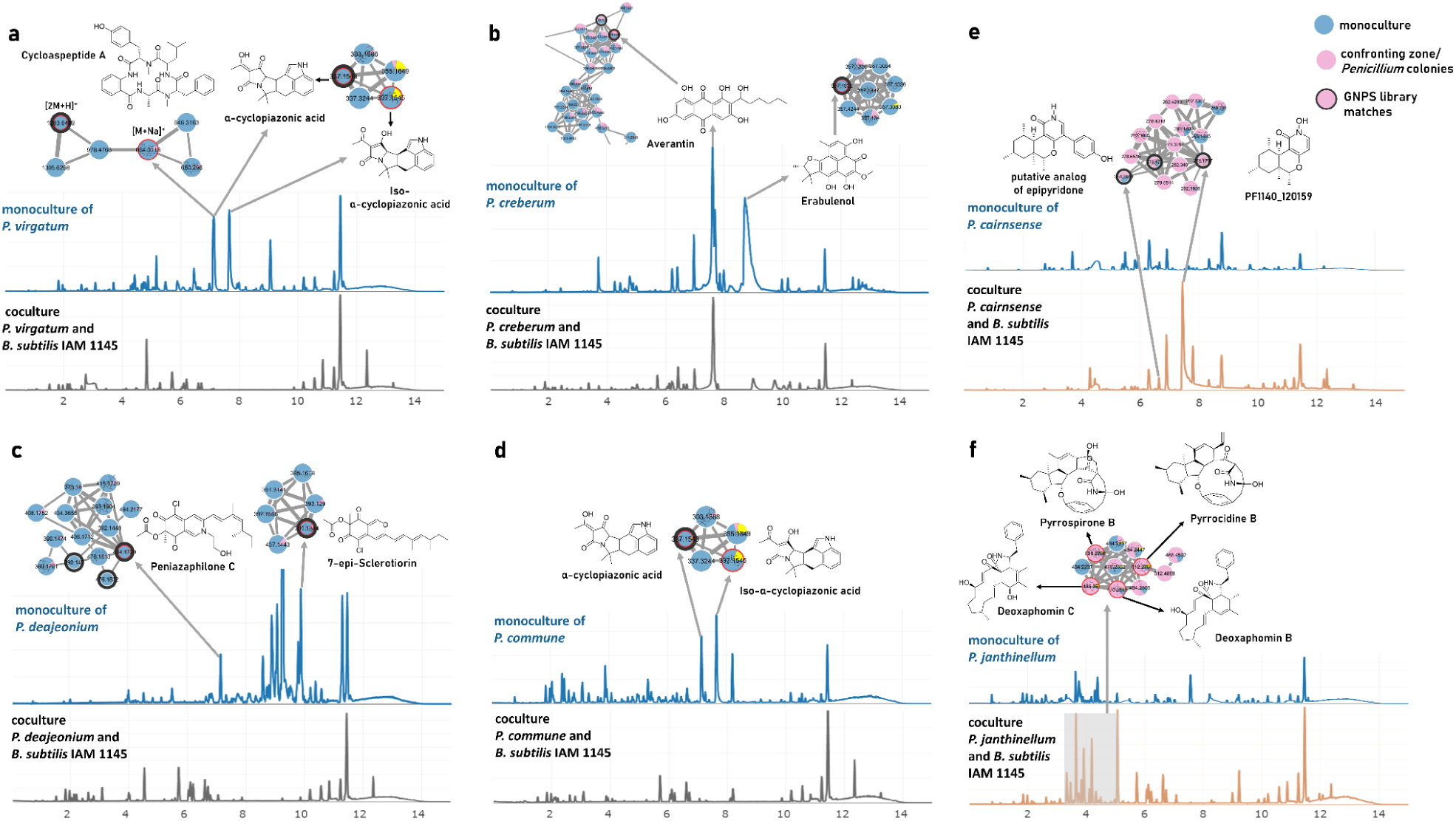
Changes in *Penicillium* metabolite profiles in cocultures with *B. subtilis* IAM 1145 based on untargeted metabolomics. BPI chromatograms of axenic and coculture conditions are shown for selected cases. Metabolite Structures were annotated via GNPS spectral matching (nodes with black borders in the MS/MS molecular network) or *in silico* annotation using NAP with the NPAtlas structural database. Specialized metaboiltes were decreased in *P. virgatum* (**a**), *P. creberum* (**b**), *P. commune* (**c**), and *P. daejeonium* compared to their monocultures. In contrast, cocultures of *P. cairnsense* (**e**) and *P. janthinellum* (**f**) showed induced production of metabolites.

**Extended Data Fig. 2.**
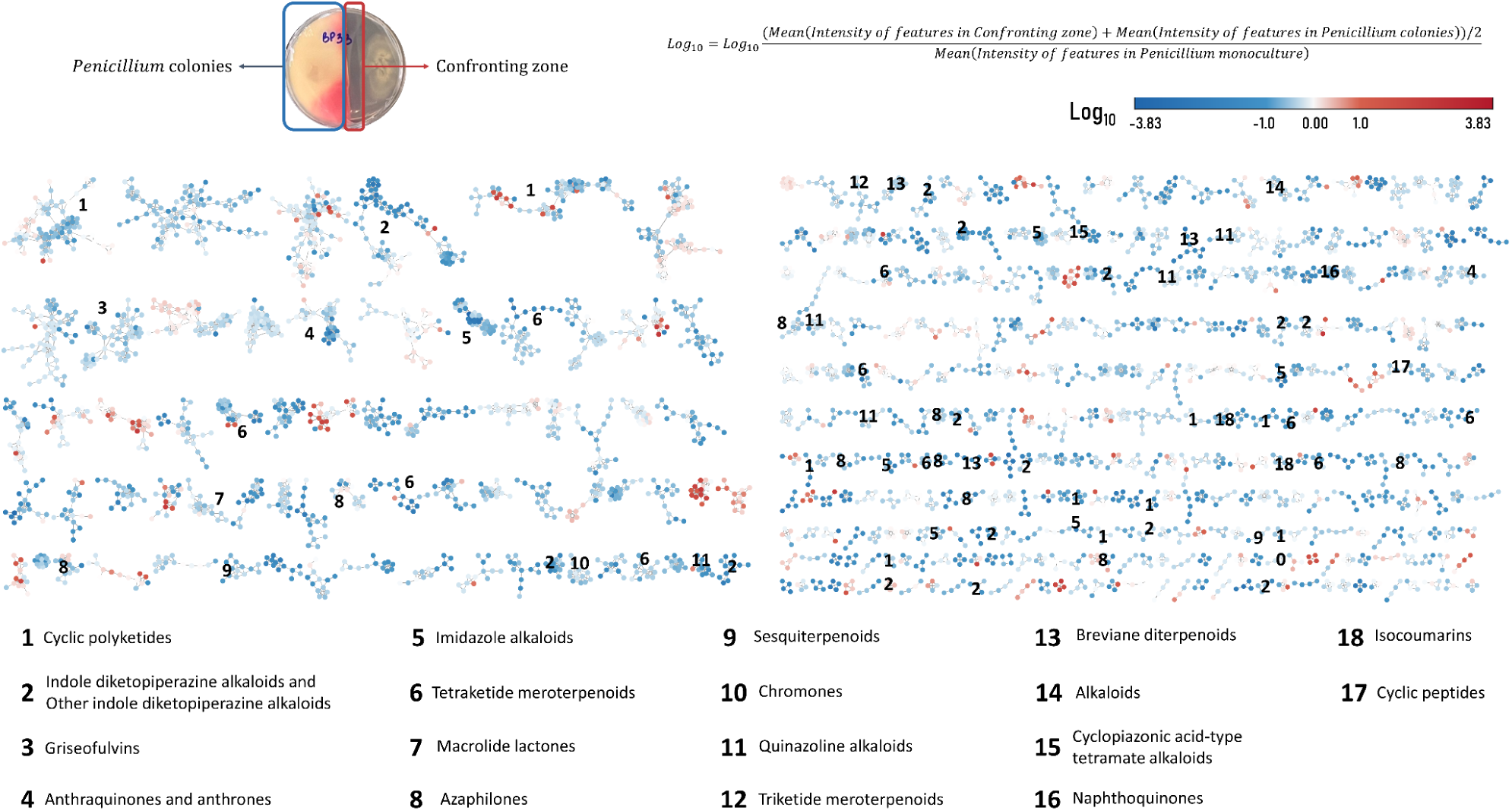
The MS/MS spectral similarity network illustrating changes in specialized metabolite levels during coculture with *B. subtilis* IAM 1145. Each node represents a single MS feature, and edges indicate MS/MS spectral similarity between features (cosine similarity ≥ 0.7). Node colors represent the log10 fold change in mean feature abundance in cocultures compared to axenic cultures.

**Extended Data Fig. 3.**
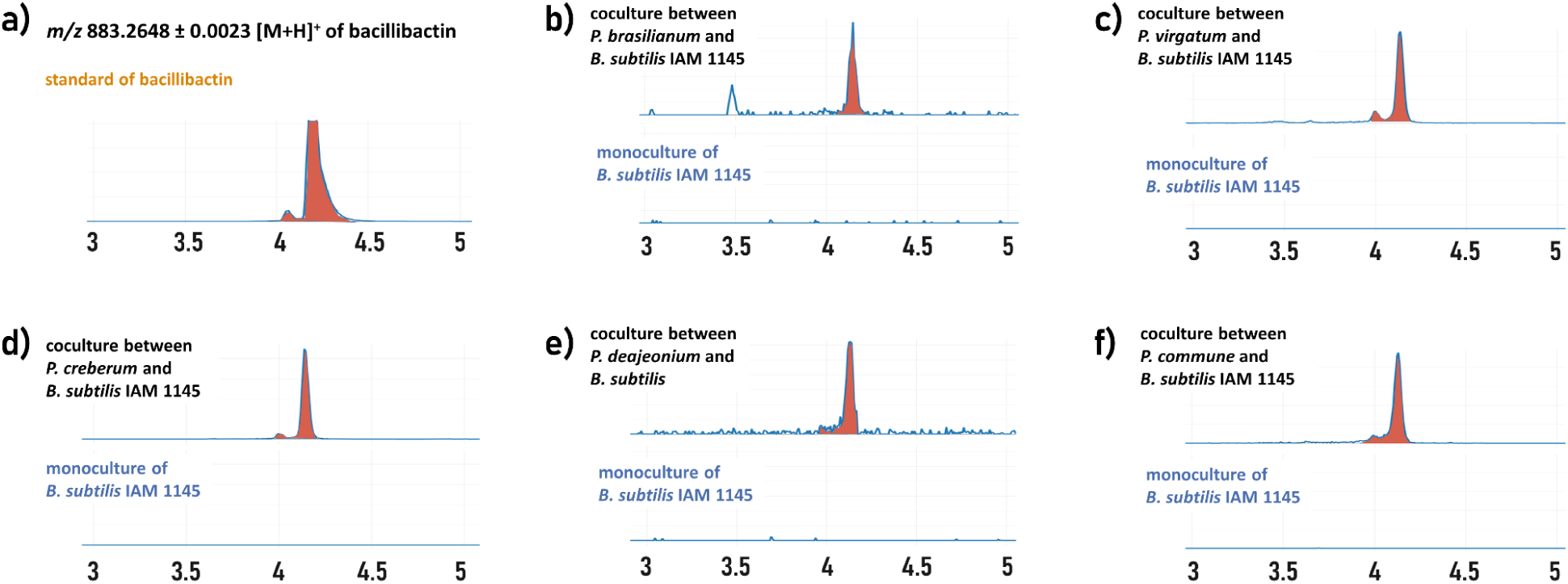
Extracted ion chromatograms (EICs) showing bacillibactin production by *B. subtilis* IAM 1145, which was induced by coculture with *Penicillium* species. (**a**) bacillibactin reference standard. (**b–f**) bacillibactin deteced in coculttures of *B. subtilis* IAM 1145 with *P. brasilianum* (**b**), *P. virgatum* (**c**), *P. creberum* (**d**), *P. deajeonium* (**e**), and *P. commune* (**f**).

**Extended Data Fig. 4.**
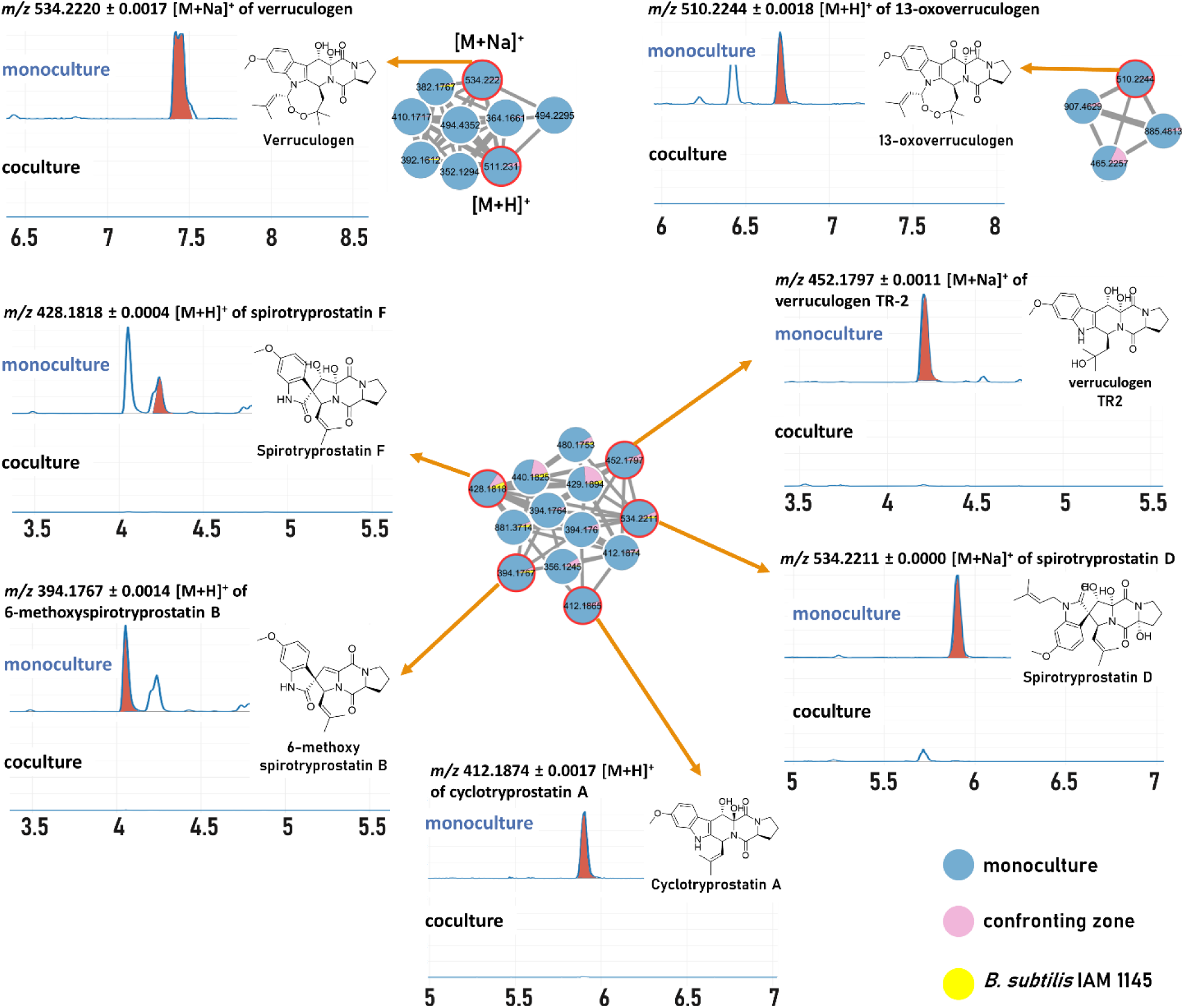
EICs of verruculogen congeners from *P. brasilianum* show decreased production in coculture with *B. subtilis* IAM 1145. Metabolites were annotated *in silico* using NAP with the NPAtlas structural library.

**Extended Data Fig. 5.**
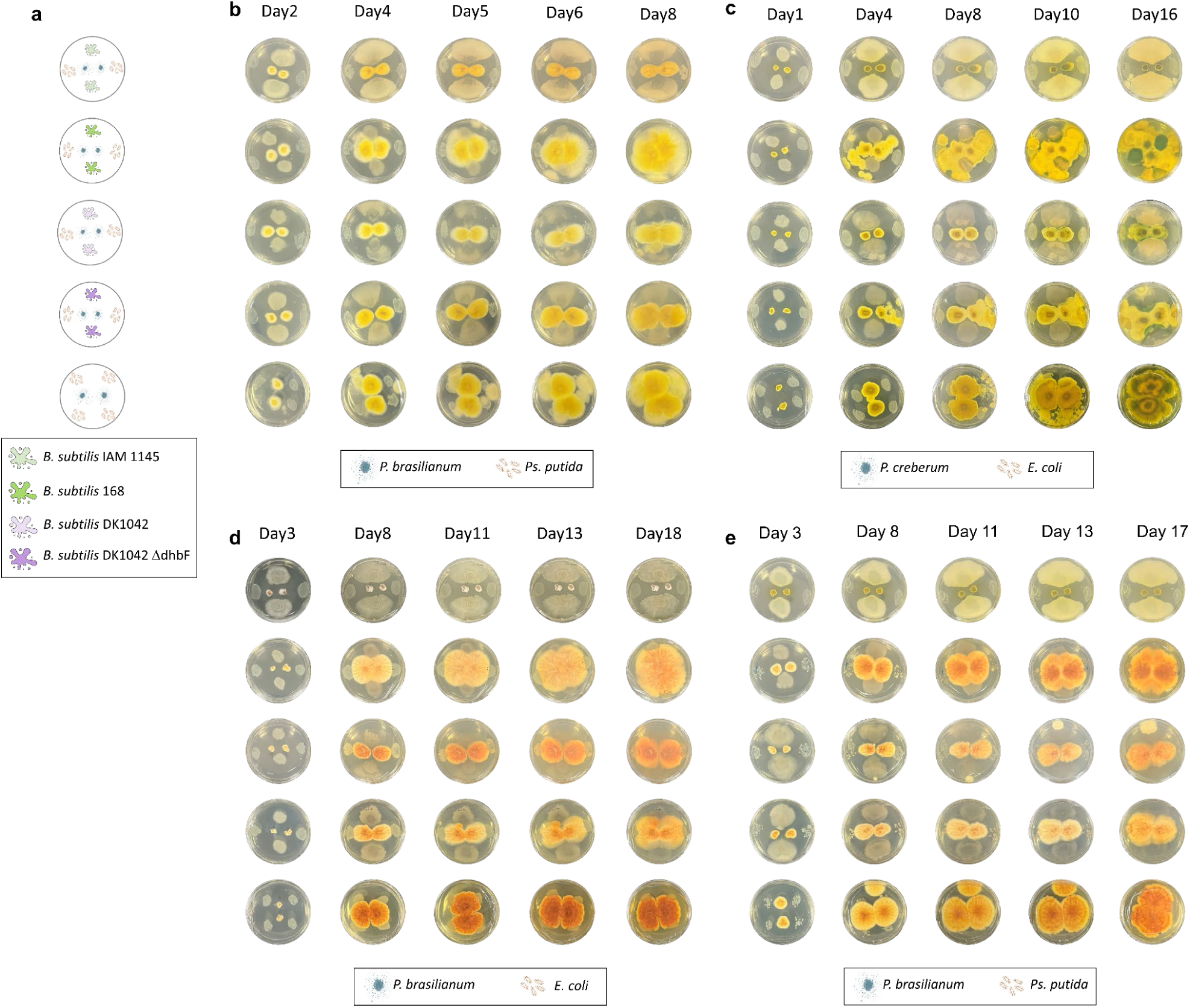
Time-course observation of tripartite cocultures involving *Penicillium* species, *B. subtilis*, and a third bacterial partner (*E. coli* or *Ps. putida*). **(a)** Schematic representation of the plate design used for the tripartite cocultures. *P. brasilianum* (**b**) and *P. virgatum* (**e**) were cultured with *Ps. putida*, while *P. creberum* (**c**) and *P. virgatum* (**d**) were cultured with *E. coli*. Each combination was monitored at multiple time points to assess the impact of bacillibactin production on fungal growth and community composition.

## References

1. Kramer, J., Özkaya, Ö. & Kümmerli, R. Bacterial siderophores in community and host interactions. Nat. Rev. Microbiol. 18, 152–163 (2020).

2. Braud, A., Geoffroy, V., Hoegy, F., Mislin, G. L. A. & Schalk, I. J. Presence of the siderophores pyoverdine and pyochelin in the extracellular medium reduces toxic metal accumulation in Pseudomonas aeruginosa and increases bacterial metal tolerance. Environ. Microbiol. Rep. 2, 419–425 (2010).

3. Schalk, I. J., Hannauer, M. & Braud, A. New roles for bacterial siderophores in metal transport and tolerance. Environ. Microbiol. 13, 2844–2854 (2011).

4. Hesse, E. et al. Ecological selection of siderophore-producing microbial taxa in response to heavy metal contamination. Ecol. Lett. 21, 117–127 (2018).

5. Achard, M. E. S. et al. An antioxidant role for catecholate siderophores in Salmonella. Biochem. J 454, 543–549 (2013).

6. Adler, C. et al. The alternative role of enterobactin as an oxidative stress protector allows Escherichia coli colony development. PLoS One 9, e84734 (2014).

7. Jin, Z. et al. Conditional privatization of a public siderophore enables Pseudomonas aeruginosa to resist cheater invasion. Nat. Commun. 9, 1383 (2018).

8. Lamont, I. L., Beare, P. A., Ochsner, U., Vasil, A. I. & Vasil, M. L. Siderophore-mediated signaling regulates virulence factor production in Pseudomonasaeruginosa. Proc. Natl. Acad. Sci. U. S. A. 99, 7072–7077 (2002).

9. Kümmerli, R. & Brown, S. P. Molecular and regulatory properties of a public good shape the evolution of cooperation. Proc. Natl. Acad. Sci. U. S. A. 107, 18921–18926 (2010).

10. Butaitė, E., Baumgartner, M., Wyder, S. & Kümmerli, R. Siderophore cheating and cheating resistance shape competition for iron in soil and freshwater Pseudomonas communities. Nat. Commun. 8, 414 (2017).

11. Sathe, S., Mathew, A., Agnoli, K., Eberl, L. & Kümmerli, R. Genetic architecture constrains exploitation of siderophore cooperation in the bacterium Burkholderia cenocepacia. Evol Lett 3, 610–622 (2019).

12. Jiricny, N. et al. Fitness correlates with the extent of cheating in a bacterium. J. Evol. Biol. 23, 738–747 (2010).

13. Schiessl, K. T., Janssen, E. M.-L., Kraemer, S. M., McNeill, K. & Ackermann, M. Magnitude and Mechanism of Siderophore-Mediated Competition at Low Iron Solubility in the Pseudomonas aeruginosa Pyochelin System. Front. Microbiol. 8, 1964 (2017).

14. Ho, Y.-N. et al. Specific inactivation of an antifungal bacterial siderophore by a fungal plant pathogen. ISME J. 15, 1858–1861 (2021).

15. Uzi-Gavrilov, S., Tik, Z., Sabti, O. & Meijler, M. M. Chemical Modification of a Bacterial Siderophore by a Competitor in Dual-Species Biofilms. Angew. Chem. Int. Ed Engl. 62, e202300585 (2023).

16. Jenul, C. et al. Pyochelin biotransformation by Staphylococcus aureus shapes bacterial competition with Pseudomonas aeruginosa in polymicrobial infections. Cell Rep. 42, (2023).

17. Andrić, S. et al. Plant-associated Bacillus mobilizes its secondary metabolites upon perception of the siderophore pyochelin produced by a Pseudomonas competitor. ISME J. 17, 263–275 (2023).

18. Charron-Lamoureux, V. et al. Pulcherriminic acid modulates iron availability and protects against oxidative stress during microbial interactions. Nat. Commun. 14, 2536 (2023).

19. Galdino, A. C. M. et al. Siderophores promote cooperative interspecies and intraspecies cross-protection against antibiotics in vitro. Nat Microbiol 9, 631–646 (2024).

20. Zang, Z. et al. Streptomyces secretes a siderophore that sensitizes competitor bacteria to phage infection. Nat. Microbiol. (2025) doi:10.1038/s41564-024-01910-8.

21. Wang, M. et al. Sharing and community curation of mass spectrometry data with Global Natural Products Social Molecular Networking. Nat. Biotechnol. 34, 828–837 (2016).

22. 22. Nothias, L. F., et al. Feature-based Molecular Networking in the GNPS Analysis Environment. *bioRxiv* 812404 (2019) doi:10.1101/812404.

23. van Santen, J. A. et al. The Natural Products Atlas: An Open Access Knowledge Base for Microbial Natural Products Discovery. ACS Cent Sci 5, 1824–1833 (2019).

24. van Santen, J. A. et al. The Natural Products Atlas 2.0: a database of microbially-derived natural products. Nucleic Acids Res. 50, D1317–D1323 (2022).

25. Sumner, L. W. et al. Proposed minimum reporting standards for chemical analysis Chemical Analysis Working Group (CAWG) Metabolomics Standards Initiative (MSI). Metabolomics 3, 211–221 (2007).

26. May, J. J., Wendrich, T. M. & Marahiel, M. A. The dhb Operon of Bacillus subtilisEncodes the Biosynthetic Template for the Catecholic Siderophore 2,3-Dihydroxybenzoate-Glycine-Threonine Trimeric Ester Bacillibactin*. J. Biol. Chem. 276, 7209–7217 (2001).

27. Fayos, J., Lokensgard, D., Clardy, J., Cole, R. J. & Kirksey, J. W. Letter: Structure of verruculogen, a tremor producing peroxide from Penicillium verruculosum. J. Am. Chem. Soc. 96, 6785–6787 (1974).

28. Kato, N. et al. Gene disruption and biochemical characterization of verruculogen synthase of Aspergillus fumigatus. Chembiochem 12, 711–714 (2011).

29. Maiya, S., Grundmann, A., Li, S.-M. & Turner, G. The fumitremorgin gene cluster of Aspergillus fumigatus: identification of a gene encoding brevianamide F synthetase. Chembiochem 7, 1062–1069 (2006).

30. Schwyn, B. & Neilands, J. B. Universal chemical assay for the detection and determination of siderophores. Anal. Biochem. 160, 47–56 (1987).

31. Pecoraro, V. L., Harris, W. R., Wong, G. B., Carrano, C. J. & Raymond, K. N. Coordination chemistry of microbial iron transport compounds. 23. Fourier transform infrared spectroscopy of ferric catechoylamide analogues of enterobactin. J. Am. Chem. Soc. 105, 4623–4633 (1983).

32. Miethke, M. et al. Ferri-bacillibactin uptake and hydrolysis in Bacillus subtilis. Mol. Microbiol. 61, 1413–1427 (2006).

33. Konkol, M. A., Blair, K. M. & Kearns, D. B. Plasmid-encoded ComI inhibits competence in the ancestral 3610 strain of Bacillus subtilis. J. Bacteriol. 195, 4085–4093 (2013).

34. Lyng, M. et al. Competition for iron shapes metabolic antagonism between Bacillus subtilis and Pseudomonas marginalis. ISME J. 18, (2024).

35. Schmeda-Hirschmann, G., Hormazabal, E., Rodriguez, J. A. & Theoduloz, C. Cycloaspeptide A and pseurotin A from the endophytic fungus Penicillium janczewskii. Z. Naturforsch. C 63, 383–388 (2008).

36. Lee, Y. M. et al. Bioactive metabolites from the sponge-derived fungus Aspergillus versicolor. Arch. Pharm. Res. 33, 231–235 (2010).

37. Asiri, I. A. M., Badr, J. M. & Youssef, D. T. A. Penicillivinacine, antimigratory diketopiperazine alkaloid from the marine-derived fungus Penicillium vinaceum. Phytochem. Lett. 13, 53–58 (2015).

38. Mourshid, S. S. A., Badr, J. M., Risinger, A. L., Mooberry, S. L. & Youssef, D. T. A. Penicilloitins A and B, new antimicrobial fatty acid esters from a marine endophytic Penicillium species. Z. Naturforsch. C 71, 387–392 (2016).

39. Saha, M. et al. Microbial siderophores and their potential applications: a review. Environ. Sci. Pollut. Res. Int. 23, 3984–3999 (2016).

40. Lee, N. et al. Iron competition triggers antibiotic biosynthesis in Streptomyces coelicolor during coculture with Myxococcus xanthus. ISME J. 14, 1111–1124 (2020).

41. Philpott, C. C. & Protchenko, O. Response to iron deprivation in Saccharomyces cerevisiae. Eukaryot. Cell 7, 20–27 (2008).

42. Shakoury-Elizeh, M. et al. Metabolic response to iron deficiency in Saccharomyces cerevisiae. J. Biol. Chem. 285, 14823–14833 (2010).

43. Jones, S. E. et al. Streptomyces volatile compounds influence exploration and microbial community dynamics by altering iron availability. MBio 10, (2019).

44. Charlang, G., Ng, B., Horowitz, N. H. & Horowitz, R. M. Cellular and extracellular siderophores of Aspergillus nidulans and Penicillium chrysogenum. Mol. Cell. Biol. 1, 94–100 (1981).

45. Hummel, W. & Diekmann, H. Preliminary characterization of ferrichrome synthetase from Aspergillus quadricinctus. Biochim. Biophys. Acta 657, 313–320 (1981).

46. Emery, T. Role of ferrichrome as a ferric ionophore in Ustilago sphaerogena. Biochemistry 10, 1483–1488 (1971).

47. Chang, C.-Y., Bajić, D., Vila, J. C. C., Estrela, S. & Sanchez, A. Emergent coexistence in multispecies microbial communities. Science 381, 343–348 (2023).

48. Chevrette, M. G. et al. Microbiome composition modulates secondary metabolism in a multispecies bacterial community. Proc. Natl. Acad. Sci. U. S. A. 119, e2212930119 (2022).

49. Gu, S. et al. Competition for iron drives phytopathogen control by natural rhizosphere microbiomes. Nat Microbiol 5, 1002–1010 (2020).

50. Cordero, O. X., Ventouras, L.-A., DeLong, E. F. & Polz, M. F. Public good dynamics drive evolution of iron acquisition strategies in natural bacterioplankton populations. Proc. Natl. Acad. Sci. U. S. A. 109, 20059–20064 (2012).

51. Koo, B.-M. et al. Construction and Analysis of Two Genome-Scale Deletion Libraries for Bacillus subtilis. Cell Syst 4, 291–305.e7 (2017).

52. Tsugawa, H. et al. MS-DIAL: data-independent MS/MS deconvolution for comprehensive metabolome analysis. Nat. Methods 12, 523–526 (2015).

53. da Silva, R. R. et al. Propagating annotations of molecular networks using in silico fragmentation. PLoS Comput. Biol. 14, e1006089 (2018).

54. Rio, D. C., Ares, M., Jr, Hannon, G. J. & Nilsen, T. W. Purification of RNA using TRIzol (TRI reagent). Cold Spring Harb. Protoc. 2010, db.prot5439 (2010).

55. Kim, D., Paggi, J. M., Park, C., Bennett, C. & Salzberg, S. L. Graph-based genome alignment and genotyping with HISAT2 and HISAT-genotype. Nat. Biotechnol. 37, 907–915 (2019).

56. Waskom, M. seaborn: statistical data visualization. J. Open Source Softw. 6, 3021 (2021).

