## Supplementary Fig. 1 for "Siderophores can alter the population dynamics of fungal-bacterial communities by inhibiting specialized metabolism"

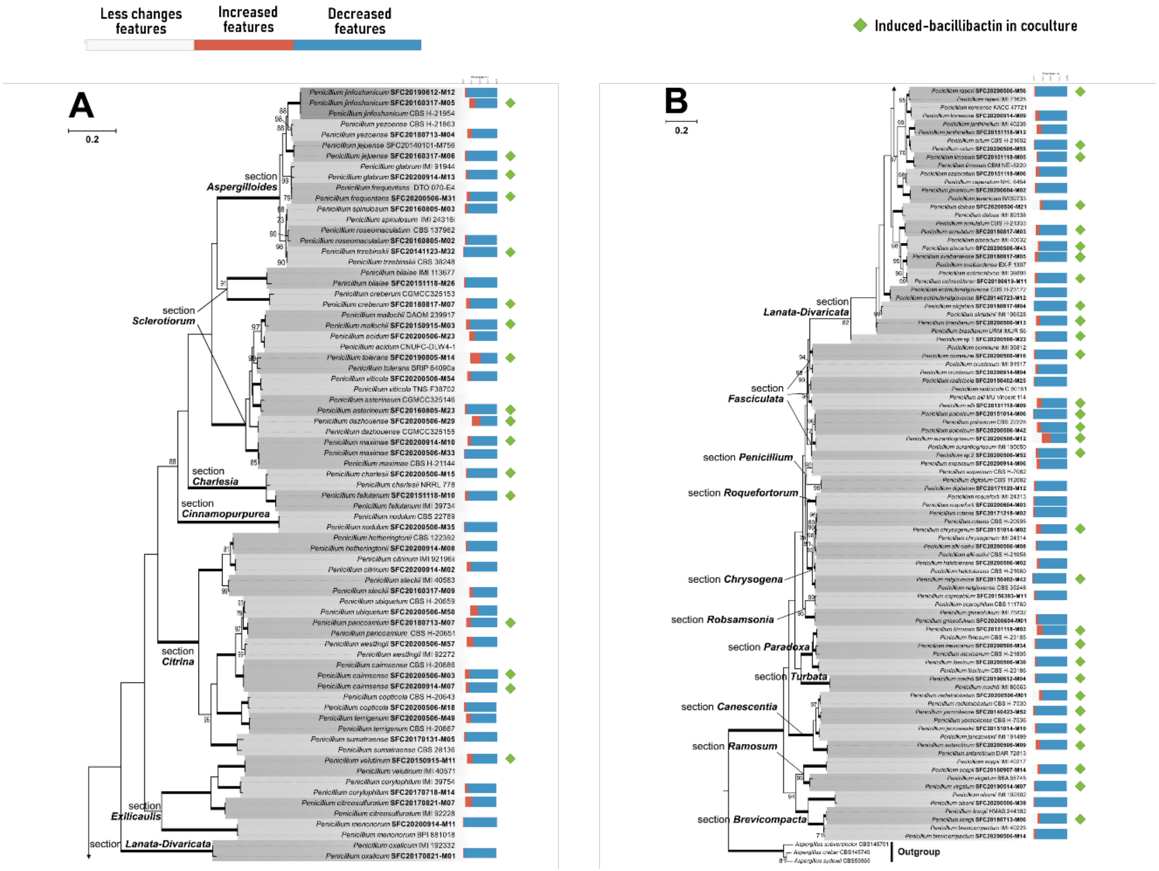


**Supplementary Fig. 1. Phylogenetic relationships and metabolite changes of *Penicillium* species in coculture with *Bacillus subtilis* IAM 1145.** The phylogenetic tree was constructed based on ITS sequences and includes sister species for reference. *Penicillium* strains used in this study are indicated in bold. Bar graphs adjacent to each strain represent the number of metabolite features that increased (upregulated) or decreased (downregulated) in coculture compared to monoculture. Green diamond symbols denote strains that induced bacillibactin production during coculture with *B. subtilis* IAM 1145**.**
