## Supplementary Fig. 2 for "Siderophores can alter the population dynamics of fungal-bacterial communities by inhibiting specialized metabolism"

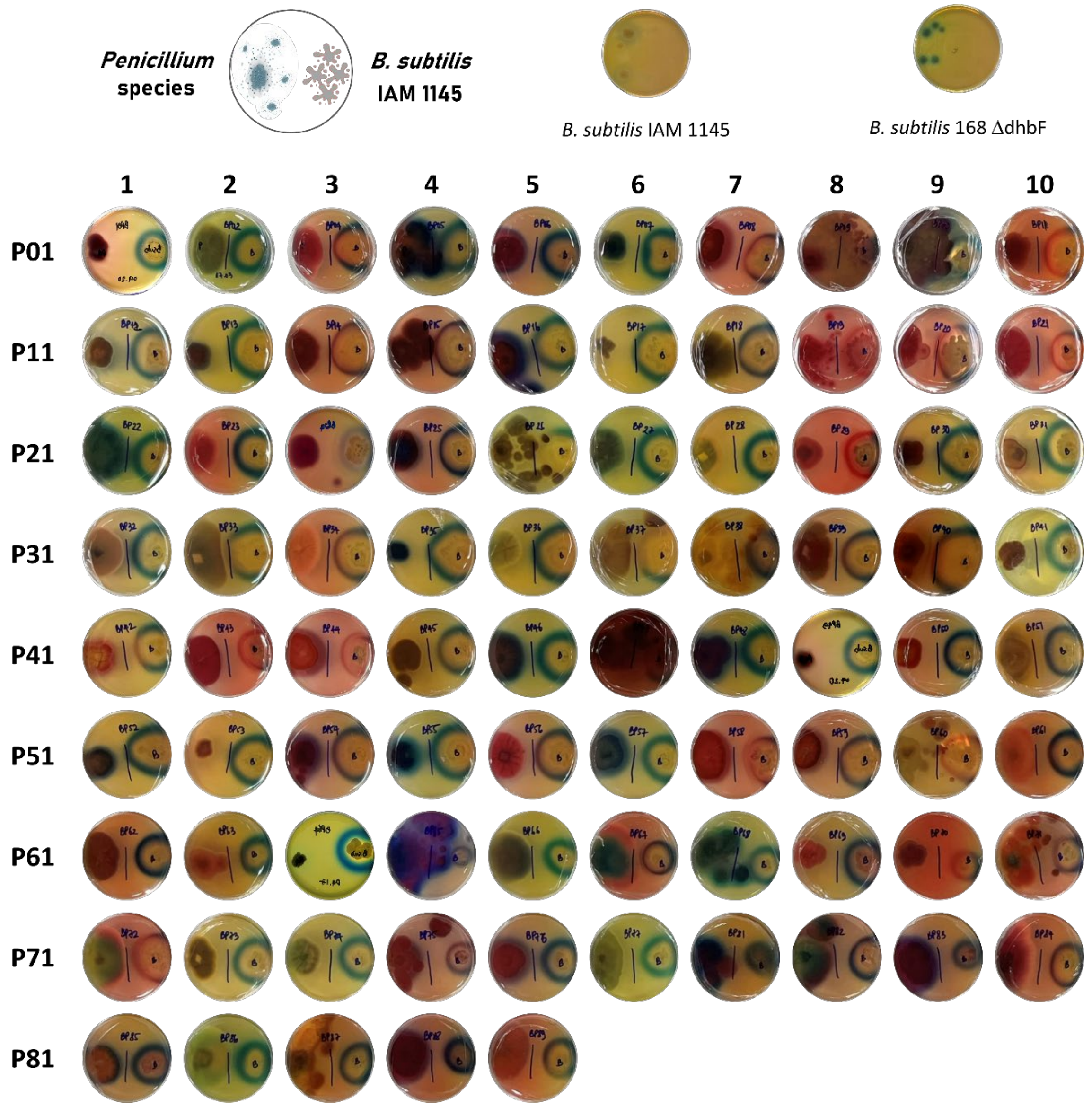


**Supplementary Fig. 2. CAS assay results of cocultures between 85 *Penicillium* species and *B. subtilis* IAM 1145 after 7 days of incubation.** A blue color indicates the presence of iron, whereas a pink color signifies iron depletion, possibly due to siderophore activity.
