## Supplementary Fig. 3 for "Siderophores can alter the population dynamics of fungal-bacterial communities by inhibiting specialized metabolism"

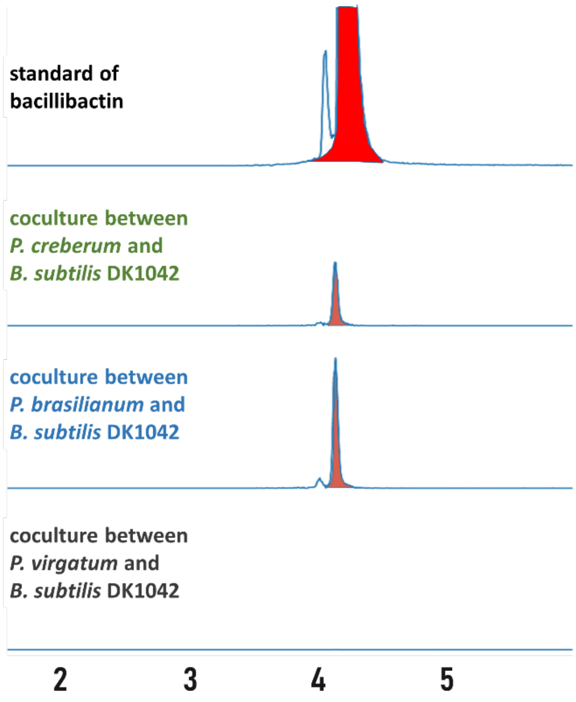


**Supplementary Fig. 3.** EICs of bacillibactin induced by *B. subtilis* DK1042 in cocultures with *P. creberum* and *P. brasilianum*. It was not induced by *P. virgatum*.
