## Supplementary Table 1 for "Siderophores can alter the population dynamics of fungal-bacterial communities by inhibiting specialized metabolism"

**Supplementary Table 1.** MS intensities of the $[M+H]^+$ feature of bacillibactin (not bound to $Fe^{3+}$) in 47 coculture plates. In the rest, bacillibactin was not detected.

| ID | Strains | Confronting zone | *Penicillium* colonies | ID | Strains | Confronting zone | *Penicillium* colonies |
|---|---|---|---|---|---|---|---|
| P01 | *Penicillium yarmokense* | 2263 | 202 | P41 | *Penicillium madriti* | 13600 | 862 |
| P03 | *Penicillium trzebinskii* | 120211 | 26533 | P43 | *Penicillium tolerans* | 850 | 0 |
| P06 | *Penicillium nalgiovense* | 0 | 335 | P44 | *Penicillium radiatolobatum* | 647 | 114 |
| P07 | *Penicillium mallochii* | 1615 | 0 | P46 | *Penicillium cairnsense* | 6182 | 1352 |
| P08 | *Penicillium velutinum* | 883 | 632 | P48 | *Penicillium antarcticum* | 306 | 0 |
| P09 | *Penicillium chrysogenum* | 103 | 647 | P49 | *Penicillium aurantiogriseum* | 0 | 1423 |
| P10 | *Penicillium polonicum* | 5054 | 1576 | P50 | *Penicillium brasilianum* | 519 | 1357 |
| P11 | *Penicillium janczewskii* | 498 | 0 | P52 | *Penicillium charlesii* | 689 | 1646 |
| P12 | *Penicillium fimosum* | 0 | 820 | P53 | *Penicillium commune* | 8052 | 5627 |
| P13 | *Penicillium limosum* | 849 | 555 | P55 | *Penicillium daleae* | 157 | 134 |
| P15 | *Penicillium allii* | 1935 | 184 | P56 | *Penicillium* sp.1 | 727 | 2137 |
| P16 | *Penicillium fellutanum* | 0 | 115 | P58 | *Penicillium dazhouense* | 4583 | 0 |
| P19 | *Penicillium jinfoshanicum* | 111 | 717 | P59 | *Penicillium ibericum* | 4229 | 1428 |
| P20 | *Penicillium jejuense* | 8855 | 0 | P60 | *Penicillium frequentans* | 3671 | 298 |
| P24 | *Penicillium asterineum* | 549 | 0 | P62 | *Penicillium mexicanum* | 496 | 1403 |
| P25 | *Penicillium soppii* | 576 | 196 | P65 | *Penicillium polonicum* | 0 | 178 |
| P32 | *Penicillium ochrochloron* | 3802 | 904 | P66 | *Penicillium piscarium* | 6175 | 1922 |
| P34 | *Penicillium kongii* | 0 | 157 | P69 | *Penicillium* sp.2 | 1931 | 867 |
| P35 | *Penicillium pancosmium* | 0 | 349 | P71 | *Penicillium ortum* | 1426 | 581 |
| P36 | *Penicillium annulatum* | 1368 | 237 | P72 | *Penicillium raperi* | 230 | 0 |
| P37 | *Penicillium skrjabinii* | 168 | 268 | P80 | *Penicillium cairnsense* | 4039 | 0 |
| P38 | *Penicillium svalbardense* | 787 | 0 | P83 | *Penicillium maximae* | 1831 | 0 |
| P39 | *Penicillium creberum* | 36597 | 176 | P85 | *Penicillium glabrum* | 467 | 1871 |
| P40 | *Penicillium virgatum* | 16316 | 0 | | | | |
