## Supplementary Table 2 for "Siderophores can alter the population dynamics of fungal-bacterial communities by inhibiting specialized metabolism"

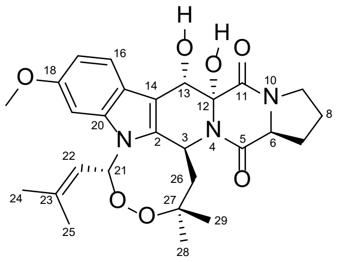

**verruculogen**

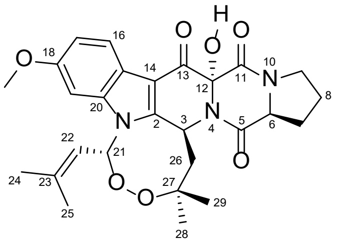

**13-oxoverruculogen**

**Supplementary Table 2.** $^{1}$H (500 MHz) and $^{13}$C (125 MHz) NMR Data of verruculogen (DMSO-$d_6$), and $^{1}$H (500 MHz) NMR Data of verruculogen and 13-oxoverruculogen ($CD_3OD$).

| | verruculogen (DMSO-$d_6$) | | verruculogen ($CD_3OD$) | 13-oxoverruculogen ($CD_3OD$) |
|---|---|---|---|---|
| **position** | $\delta_C$ | $\delta_H$ (*J* in Hz) | $\delta_H$ (*J* in Hz) | $\delta_H$ (*J* in Hz) |
| **2** | 130.9 | | | |
| **3** | 47.8 | 5.87, d (10.1) | 6.07, d (10.3) | 6.31, dd (9.8, 1.2) |
| **5** | 165.9 | | | |
| **6** | 58.4 | 4.45, dd (9.2, 6.9) | 4.54, dd (9.0, 7.3) | 4.81, dd (9.1, 7.6) |
| **7** | 22.1 | 1.94, m | 1.99, m | 1.92, dd (14.0, 1.3) |
| | | | 1.67, m | 1.82, dd (14.0, 1.3) |
| **8** | 28.9 | 2.31, m | 2.46, m | 2.39, m |
| | | 1.94, m | 1.99, m | 1.97, m |
| **9** | 45.1 | 3.46, m | 3.61, m | 3.49, m |
| **11** | 170.6 | | | |
| **12** | 82.7 | | | |
| **13** | 67.9 | 5.41, dd (3.4, 1.1) | 5.57, d (0.8) | - |
| **14** | 107.3 | | | |
| **15** | 135.8 | | | |
| **16** | 121.3 | 7.73, d (8.7) | 7.84, d (8.7) | 8.10, d (8.7) |
| **17** | 108.9 | 6.70, dd (8.7, 2.3) | 6.71, m | 6.95, dd (8.7, 2.2) |
| **18** | 155.5 | | | |
| **19** | 93.7 | 6.76, d (2.3) | 6.64, d (2.1) | 6.80, d (2.1) |
| **20** | 120.7 | | | |
| **21** | 85.1 | 6.82, d (8.2) | 6.73, m | 6.89, d (8.0) |
| **22** | 118.4 | 5.01, dt (8.3, 1.4) | 5.11, m | 5.00, dp (8.0, 1.4) |
| **23** | 142.6 | | | |
| **24** | 18.5 | 1.98, d (1.4) | 2.03, d (1.1) | 2.04, d (1.4) |
| **25** | 25.2 | 1.7, d (1.4) | 1.70, d (1.1) | 1.76, d (1.4) |
| **26** | 50.8 | 1.89, m | 2.08, m | 2.14, m |
| | | 1.58, m | | 1.94, m |
| **27** | 81.6 | | | |
| **28** | 24.1 | 1.58, s | 1.66, s | 1.63, s |
| **29** | 26.7 | 0.96, s | 1.00, s | 1.00, s |
| **12-OH** | | 6.47, s | | |
| **13-OH** | | 5.27, s | | |
| **O-CH3** | 55.2 | 3.76, s | 3.81, s | 3.82, s |
