## Supplementary Table 3 for "Siderophores can alter the population dynamics of fungal-bacterial communities by inhibiting specialized metabolism"

**Supplementary Table 3.** The list of *Penicillium* species obtained from the Korea Marine Fungi Resource Bank.

| ID | Strain code | Identification | ID | Strain code | Identification |
|---|---|---|---|---|---|
| **P01** | SFC20140423-M52 | *Penicillium yarmokense* | **P44** | SFC20200506-M01 | *Penicillium radiatolobatum* |
| **P02** | SFC20140723-M12 | *Penicillium echinulonalgiovense* | **P45** | SFC20200506-M02 | *Penicillium halotolerans* |
| **P03** | SFC20141123-M32 | *Penicillium trzebinskii* | **P46** | SFC20200506-M03 | *Penicillium cairnsense* |
| **P04** | SFC20150303-M11 | *Penicillium coprophilum* | **P47** | SFC20200506-M08 | *Penicillium allii-sativi* |
| **P05** | SFC20150402-M25 | *Penicillium radicicola* | **P48** | SFC20200506-M09 | *Penicillium antarcticum* |
| **P06** | SFC20150402-M42 | *Penicillium nalgiovense* | **P49** | SFC20200506-M12 | *Penicillium aurantiogriseum* |
| **P07** | SFC20150915-M03 | *Penicillium mallochii* | **P50** | SFC20200506-M13 | *Penicillium brasilianum* |
| **P08** | SFC20150915-M11 | *Penicillium velutinum* | **P51** | SFC20200506-M14 | *Penicillium brevicompactum* |
| **P09** | SFC20151014-M02 | *Penicillium chrysogenum* | **P52** | SFC20200506-M15 | *Penicillium charlesii* |
| **P10** | SFC20151014-M06 | *Penicillium polonicum* | **P53** | SFC20200506-M16 | *Penicillium commune* |
| **P11** | SFC20151014-M10 | *Penicillium janczewskii* | **P54** | SFC20200506-M18 | *Penicillium copticola* |
| **P12** | SFC20151118-M03 | *Penicillium fimosum* | **P55** | SFC20200506-M21 | *Penicillium daleae* |
| **P13** | SFC20151118-M05 | *Penicillium limosum* | **P56** | SFC20200506-M22 | *Penicillium* sp.1 |
| **P14** | SFC20151118-M06 | *Penicillium caperatum* | **P57** | SFC20200506-M23 | *Penicillium acidum* |
| **P15** | SFC20151118-M09 | *Penicillium allii* | **P58** | SFC20200506-M29 | *Penicillium dazhouense* |
| **P16** | SFC20151118-M10 | *Penicillium fellutanum* | **P59** | SFC20200506-M30 | *Penicillium ibericum* |
| **P17** | SFC20151118-M12 | *Penicillium janthinellum* | **P60** | SFC20200506-M31 | *Penicillium frequentans* |
| **P18** | SFC20151118-M26 | *Penicillium bilaiae* | **P61** | SFC20200506-M33 | *Penicillium maximae* |
| **P19** | SFC20160317-M05 | *Penicillium jinfoshanicum* | **P62** | SFC20200506-M34 | *Penicillium mexicanum* |
| **P20** | SFC20160317-M06 | *Penicillium jejuense* | **P63** | SFC20200506-M35 | *Penicillium nodulum* |
| **P21** | SFC20160317-M09 | *Penicillium steckii* | **P64** | SFC20200506-M39 | *Penicillium olsonii* |
| **P22** | SFC20160805-M02 | *Penicillium roseomaculatum* | **P65** | SFC20200506-M42 | *Penicillium polonicum* |
| **P23** | SFC20160805-M03 | *Penicillium spinulosum* | **P66** | SFC20200506-M43 | *Penicillium piscarium* |
| **P24** | SFC20160805-M23 | *Penicillium asterineum* | **P67** | SFC20200506-M49 | *Penicillium terrigenum* |
| **P25** | SFC20160907-M14 | *Penicillium soppii* | **P68** | SFC20200506-M50 | *Penicillium ubiquetum* |
| **P26** | SFC20170131-M05 | *Penicillium sumatraense* | **P69** | SFC20200506-M52 | *Penicillium* sp.2 |
| **P27** | SFC20170718-M14 | *Penicillium corylophilum* | **P70** | SFC20200506-M54 | *Penicillium viticola* |
| **P28** | SFC20170821-M01 | *Penicillium oxalicum* | **P71** | SFC20200506-M55 | *Penicillium ortum* |
| **P29** | SFC20170821-M07 | *Penicillium citreosulfuratum* | **P72** | SFC20200506-M56 | *Penicillium raperi* |
| **P30** | SFC20171120-M12 | *Penicillium digitatum* | **P73** | SFC20200506-M57 | *Penicillium westlingii* |
| **P31** | SFC20171218-M02 | *Penicillium rubens* | **P74** | SFC20200604-M01 | *Penicillium griseofulvum* |
| **P32** | SFC20180619-M11 | *Penicillium ochrochloron* | **P75** | SFC20200604-M02 | *Penicillium javanicum* |
| **P33** | SFC20180713-M04 | *Penicillium yezoense* | **P76** | SFC20200604-M03 | *Penicillium roqueforti* |
| **P34** | SFC20180713-M06 | *Penicillium kongii* | **P77** | SFC20200914-M02 | *Penicillium citrinum* |
| **P35** | SFC20180713-M07 | *Penicillium pancosmium* | **P78** | SFC20200914-M04 | *Penicillium crustosum* |
| **P36** | SFC20180817-M03 | *Penicillium annulatum* | **P79** | SFC20200914-M06 | *Penicillium expansum* |
| **P37** | SFC20180817-M04 | *Penicillium skrjabinii* | **P80** | SFC20200914-M07 | *Penicillium cairnsense* |
| **P38** | SFC20180817-M05 | *Penicillium svalbardense* | **P81** | SFC20200914-M08 | *Penicillium hetheringtonii* |
| **P39** | SFC20180817-M07 | *Penicillium creberum* | **P82** | SFC20200914-M09 | *Penicillium koreense* |
| **P40** | SFC20190514-M07 | *Penicillium virgatum* | **P83** | SFC20200914-M10 | *Penicillium maximae* |
| **P41** | SFC20190612-M04 | *Penicillium madriti* | **P84** | SFC20200914-M11 | *Penicillium menonorum* |
| **P42** | SFC20190612-M12 | *Penicillium jinfoshanicum* | **P85** | SFC20200914-M13 | *Penicillium glabrum* |
