## Supplementary Table 4 for "Siderophores can alter the population dynamics of fungal-bacterial communities by inhibiting specialized metabolism"

**Supplementary Table 4.** The composition of the defined iron-deficient medium.

| **Components** | **Defined iron-deficient** | **Defined iron-deficient + iron** |
|---|---|---|
| Glucose | 10.0 g | 10.0 g |
| Peptone | 2.0 g | 2.0 g |
| $MgSO_4.7H_2O$ | 0.5 g | 0.5 g |
| $KH_2PO_4$ | 1.0 g | 1.0 g |
| Agar | 15.0 g | 15.0 g |
| $FeNH_4(SO_4)_2.12H_2O$ | - | 100 µM |
| Distilled water | 1000 mL | 1000 mL |
